# Signatures of Electron-hole Hopping in Myoglobin Peroxidase Activity Revealed by Deep Mutational Learning

**DOI:** 10.1101/2025.08.27.672588

**Authors:** Christoph Küng, Alperen Dalkıran, Rosario Vanella, Diego A. Oyarzún, Michael A. Nash

## Abstract

In addition to storing molecular oxygen, myoglobin catalyzes peroxidase-like reactions involving high valency iron(IV)-oxo species that support one-electron oxidations on a range of substrates at an open active site. In select metalloenzymes, long-range electron transfer can be mediated by hole-hopping pathways composed of aromatic residues that act as relay stations for oxidative equivalents. However, it remains unclear how sequence variations could introduce or alter such catalytic mechanisms in myoglobin. Here we used enzyme proximity sequencing (EP-Seq) to measure the peroxidase activity levels of >6,000 human myoglobin variants. The resulting fitness landscape reveals how aromatic substitutions, in particular surface-exposed tryptophans, can enhance peroxidase activity. Using protein language models in tandem with feedforward neural networks, we trained an accurate fitness predictor on the experimental dataset, and applied it to screen >4M double mutant variants. The predictions suggested a beneficial role for electron-hole-hopping mutations in improving peroxidase activity. We experimentally tested 20 high scoring variants in a yeast display assay, all of which outperformed wild type myoglobin. Three selected variants were also tested in soluble format and similarly showed improved performance. A focused combinatorial library yielded a top double tryptophan variant (Q92W/F107W) with 4.9-fold higher catalytic efficiency than wild type. These results show that deep mutational learning can identify myoglobin variants with enhanced peroxidase activity that are consistent with the involvement of hole-hopping pathways, with broad implications for biocatalyst and redox enzyme design.

## Introduction

Myoglobin is a small, globular heme protein found predominantly in the cardiac and skeletal muscle of vertebrates, comprising a single polypeptide chain folded into eight α-helices^1^. Best known for its role in oxygen storage and transport, myoglobin buffers oxygen levels during high metabolic demand.^2^ In addition to this classical function, myoglobin has more recently attracted interest for its catalytic activity^3,4^. This arises from its iron-containing protoporphyrin IX cofactor, which can reversibly cycle oxidation states to catalyze redox reactions. In the presence of hydrogen peroxide (H_2_O_2_), the iron center can be oxidized to form high-valent iron(IV)-oxo species referred to as Compound I/II. These reactive intermediates can catalyze one-electron oxidations on a broad range of organic substrates facilitated by an open and accessible active site geometry.

The peroxidase activity of myoglobin is normally suppressed in myocytes due to the reducing environment of the cell. However, there is evidence that this activity can occur *in vivo* under pathological conditions^5–7^. For example, in rhabdomyolysis or upon reperfusion of ischemic tissue, increased levels of reactive oxidative species (ROS) create conditions that support heme-mediated oxidation.^8–10^ Myoglobin peroxidase activity may contribute to ROS detoxification in these settings, but it can also lead to oxidative damage^9,11^. When endogenous antioxidants like ascorbate or glutathione are depleted, myoglobin can oxidize lipids and damage proteins and DNA. Recognizing this expanded catalytic potential, researchers have sought to engineer myoglobin for a range of applications, including dye decolorization, and antibiotic degradation.^12–16^

Multiple studies point to tyrosine and tryptophan substitutions as important for engineering electron transfer pathways in peroxidases. These aromatic residues either directly enhance peroxidase activity, or increase cofactor reduction rates by electron donors. Their evolutionary appearance has been linked to the oxygenation of Earth’s atmosphere, suggesting an adaptive response to oxidative stress^17–20^. Prior work by Gray and Winkler has emphasized both the catalytic and protective roles of such residues in natural enzymes, showing that they can extend redox activity beyond the active site and help shuttle oxidative equivalents through protein scaffolds.^21–23^ Introducing these residues into a stable single-domain protein like myoglobin offers a powerful platform for dissecting protein-based radical chemistry and guiding rational enzyme design.

To systematically understand how mutations could influence the peroxidase-like activity of myoglobin, we sought out a high-throughput method that could provide suitable data for variant discovery with machine learning (ML). Deep mutational scanning (DMS) is one such powerful approach that enables massively parallel analysis of protein sequence-function pairings.^24–26^ However, building high-throughput platforms to assay enzymatic activity remains challenging.^27^ Linking genotype to enzymatic phenotype requires the ability to compartmentalize enzymatic reactions, as well as distinguish between signal generation due to improved catalytic properties from that attributable to higher enzyme abundance (i.e. expression level). Although droplet-based systems with colorimetric readouts^28–30^ and survival-based selections^31–33^ can be used to enrich functional variants, these approaches suffer from infrastructure and biochemical limitations, and frequently confound improved enzyme activity with increased protein expression levels.

To address this challenge, we recently developed enzyme proximity sequencing (EP-Seq), a high-throughput method based on yeast display that enables parallel assessment of both protein expression levels and enzymatic activity (**Figure 1**)^26,34,35^. This platform uses phenoxy radical-based labelling chemistry to couple enzymatic activity to yeast cell fluorescence. Highly active variants can then be separated from inactive variants by fluorescence-activated cell sorting (FACS). In prior work, this strategy relied on exogenous horseradish peroxidase for radical labeling, however in the present study, we used the intrinsic peroxidase activity of surface-displayed myoglobin^34^. Leveraging our high-throughput dataset, we then applied machine learning to model the mutational fitness landscape of human myoglobin and predict peroxidase activity of all double mutants.

**Figure 1).**
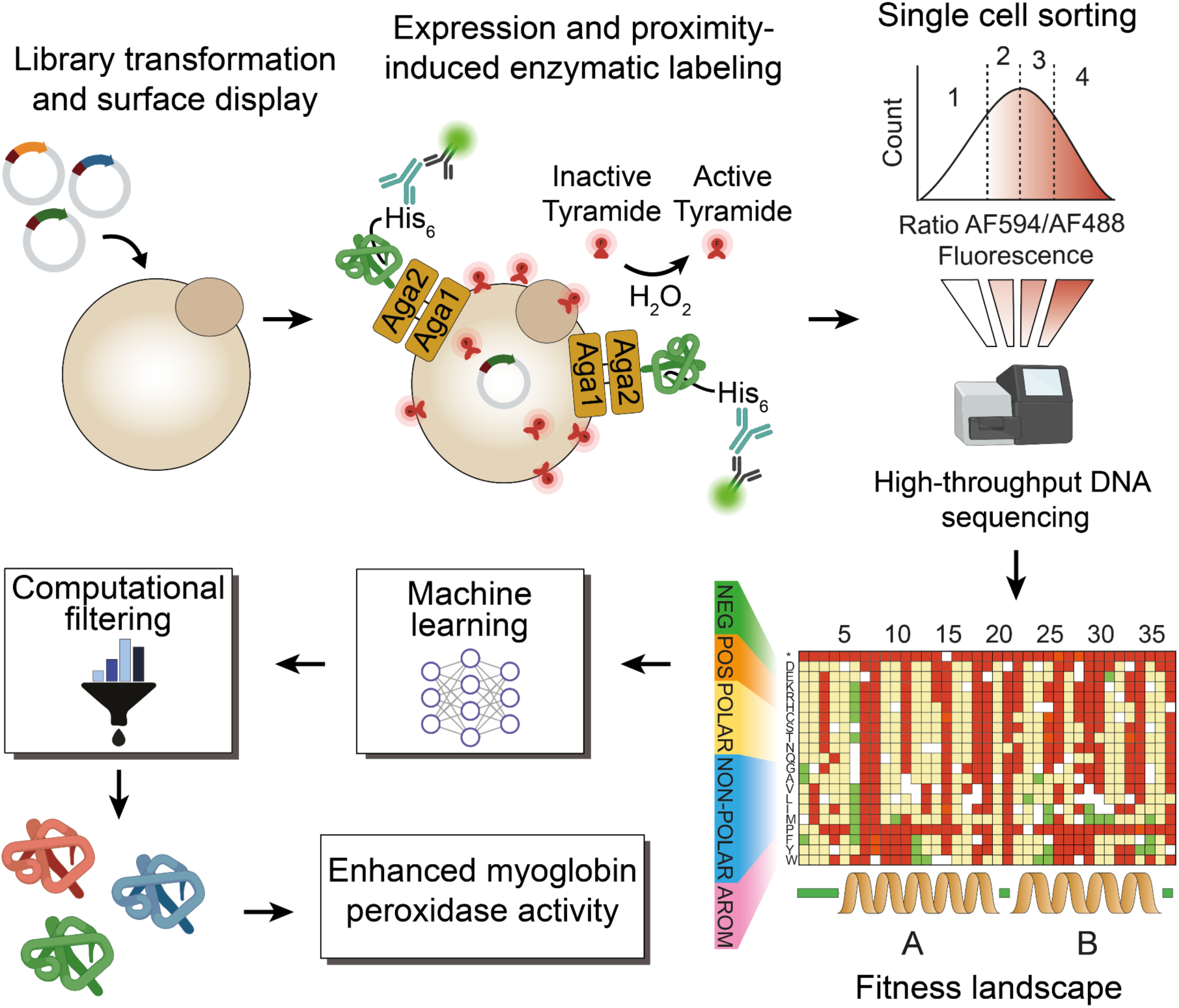
Experimental scheme. A barcoded library is transformed in EBY100 (*S. Cerevisiae*) strain and encoded protein variants are displayed on the cell surface. After immunostaining to quantify expression, the peroxidase activity of myoglobin is used for enzyme-mediated radical proximity labelling *via* reaction with the peroxidase substrate tyramide-Alexa Fluor 594. This labels the single cells in proportion to the peroxidase activity level of the myoglobin variants display on its surface, allowing library screening in a one-pot reaction. The 2-color labeled library is then sorted into four bins by fluorescent activated cell sorting (FACS) based on the expression-normalized peroxidase activity level. Variant distribution amongst bins is evaluated by next generation sequencing and converted into fitness scores. This largescale fitness landscape is then used to train a variety of machine learning algorithms to predict peroxidase activity from sequence. After model training, *in silico* screening and computational filtering of the highest ranked hits is undertaken to arrive at a final subset which is characterized biochemically in a soluble format. What emerges from this pipeline are highly ranked variants featuring surface-accessible Tyr and Trp substitutions that implicate hole-hopping–mediated access of oxidative equivalents to the active site and improved catalytic activity.

Recent advances have established ML as a powerful tool for protein variant prediction^36^. The fusion of deep mutational scanning with machine learning, termed deep mutational learning (DML), offers an efficient method to gain deeper understanding in sequence-function relationships^37,38^. Unlike many ML approaches that yield opaque predictions, our DML framework facilitated interpretable hypothesis generation by recommending substitutions with residues that enable hole hopping. This connection, learned from the training data, illustrates how integrating DMS with ML can yield mechanistic insights rather than serving solely as an uninterpretable prediction tool.

This analysis led to the identification of highly active variants which were validated experimentally. Importantly, we further demonstrated that activity trends observed in the yeast-display format were recapitulated in the corresponding soluble enzyme versions, underscoring the generalizability of our approach.

## Materials & Methods

### Materials

Restriction enzymes, DNA ligase kit and calf intestinal phosphatase (CIP) were purchased from New England Biolabs (Beverly, MA). GelRed^TM^ dye nucleic acid gel stain was purchased from Biotium. Gene Morph II random mutagenesis kit (epPCR) was obtained from Agilent Technologies (Santa Clara, CA). The primary and secondary antibodies were purchased from Thermo Fisher Scientific (Waltham, MA). Custom oligonucleotides were obtained from Microsynth (Balgach, CH). The DNA miniprep, gel purification and PCR purification kits were purchased from Thermo Fisher Scientific. Chemicals, if not indicated otherwise, are from Sigma Aldrich (St. Louis, MO). Sodium alginate was purchased from Duchefa Biochemie (Haarlem, Netherlands).

### Myoglobin gene cloning

The homo sapiens myoglobin (Mb) gene was obtained from TWIST Bioscience codon optimized for *Saccharomyces Cerevisiae*. The gene was cleaved in a double digest with restriction enzymes EcoRI and XhoI and cloned in a pYD1 expression vector. Sequence was confirmed by sanger sequencing and the plasmid featuring the myoglobin gene transformed in yeast strain EBY100 by a lithium acetate transformation protocol ^39^. Positive colonies were selected on synthetic defined (SD) agar plates lacking tryptophan (-TRP) supplemented with glucose 2% (wt/vol) and ampicillin.

### Construction of barcoded myoglobin mutant library

For mutagenesis of the human myoglobin wild type gene, we used a plasmid based one-pot mutagenesis protocol which was previously described ^40^. Since higher efficiency was observed for smaller plasmids, we transferred the expression cassette from the pYD1 (5446 bp) to a smaller puc19 plasmid (3647 bp). The puc19 plasmid contains two BbvCI restriction sites up- and downstream of the expression cassette. The original protocol was slightly modified by adding an extra step after the first incubation with Nt.BbvCI enzyme and exonuclease. The reaction mixture was followed by incubation with 10 units of quick CIP phosphatase (New England Biolabs) at 37°C for 20 min, with subsequent incubation at 80°C for 20 minutes. With this step, the removal of 5’ phosphate from nicked and partially degraded strands of wild type gene is promoted and hence limits the reformation of closed wild type plasmids in the following amplification and ligation steps. Then, nested NNK codon containing primers were used for synthesis of the mutated strand targeting codon triplets 1 to 154 (full gene) of the myoglobin wild type gene. Primers were designed with the script create_primers.py from ^40^ whereas the length of the primers was adjusted to reach a similar melting temperature around 60 °C for all primers. Designed primers were ordered from Integrated DNA technologies in a 96 well plate format and mixed in equimolar ratios to the final concentration of 10 µM. With the mixed primers, the site saturation protocol was performed as described in ^40^. After obtaining the plasmid based mutation library, molecular barcodes were introduced through PCR and hence linked to each of the amplified Myoglobin variants. PCR was accomplished by using primers P1 and P2 shown in **Table S1**, adding a 15 N barcode in front of the galactose promoter sequence. The amplified and barcoded library was cloned in a pYDKan backbone, by HiFi DNA Assembly (New England Biolabs) and using the vector plasmid which was linearized by BbVCI/PmeI double digestion. After column purification (Zymo-Spin I, Zymoresearch), the HiFi assembly product was then transformed into electrocompetent XL1-Blue (Agilent) cells by electroporation (Gene Pulser, Biorad).

Successfully transformed cells were selected for in LB agar plates supplemented with (50 μg/ml) Kanamycin and the size of the library was estimated by counting colony forming units on plates with serially diluted fractions of the library. The total size of the barcoded library was estimated to be 300,000 variants. After incubation at 37°C for 20 hours, the colonies were scraped from the plates and resuspended in liquid LB Kan, further grown for 2 hours and then plasmids extracted by standard Miniprep method. 10 μg from the plasmid library pool was introduced into the *Saccharomyces cerevisiae* strain EBY100, using the lithium acetate transformation protocol^39^. To assess the efficiency of the transformation, successive dilutions of the transformed material were spread onto agar plates containing -Trp and 2% (wt/vol) glucose. Counting of the resulting colony forming units revealed a transformation efficiency around 2 million colonies. Following the transformation, the yeast mutant library was cultivated for 24 hours at a temperature of 30°C in a liquid medium lacking tryptophan (-Trp), supplemented with 2% (wt/vol) glucose. Subsequently, the culture was diluted to an OD_600_ of 1 and transferred to fresh -Trp medium with 2% (wt/vol) glucose to minimize the chance of obtaining multiple transformants. This culture was then allowed to grow for an additional 16 hours at 30°C. Finally, portions containing approximately 5 *10^7 cells were prepared and suspended in a solution of 25% (vol/vol) glycerol before being preserved at -80°C. Plasmid maps are provided at: https://zenodo.org/uploads/16781055.

In order to limit the size of the library and increase the confidence in barcode assignment from the PacBio-generated lookup table, we capped/bottlenecked the library to 100,000 cells. This was done on a FACS Sony SH800S Sorter by simply gating for viable single cells and sorting 100,000 events in growth media. This culture was grown for 20 h overnight and stored as glycerol stocks at -80°C. The capping was done in yeast cells, since better survival rate was observed when compared to *E.Coli*.

### PacBio long read sequencing

In order to link the molecular barcodes to their respective hMb variant sequence, Pacbio long read sequencing was used. The capped library glycerol stock was thawed and used to inoculate 40 mL -TRP and 2% (wt/vol) glucose at an initial OD of 0.1 and allowed to grow for 24 hours. OD then was around 6.8 and 1 mL (6 samples) of the cultures were collected, spun down, and resuspended in 250 µl of Miniprep Resuspension buffer 1 (GeneJET, ThermoFisher). Then 4 µl of Zymolyase was added (Zymoresearch) and incubated for 2 h at 37°C with shaking at 1,000 rpm. Then the standard protocol as suggested in the kit was followed, whereas the final elution volume was 10 µl of prewarmed nuclease free sterile water. Then, the capped library was transformed into electrocompetent E.Coli by electroporation of 6 individual reactions. After outgrowth, amplified plasmids were purified by miniprep and 8 ug were digested by restriction enzymes Not1 and Pme1 for 16 hours. The fragment of interest was isolated by agarose gel electrophoresis and after gel purification of corresponding bands, DNA was cleaned by AMPure XP bead purification.

The purified DNA was then used for SMRT bell ligation and run on a Pacific Biosciences Sequel IIa sequencer with a movie time of 15 hours. Resulting sequences were filtered for quality score Phred 20 and higher. Additionally, sequences were filtered for a maximal length of 1800 bp (amplicon length is 1569). Then, the computational workflow already established and reported in ^26^ was used to extract the barcode and assign it to a respective mutation. We used Minimap2 ^41^ to map the long read sequences to a reference file with the wild type sequence of the expression vector PCR amplicon. Then, a C script was used to process the same file output of the mapped sequences using a custom library file (libscodon.h). The script extracts the 15 base pair molecular barcode and its quality along with the mutational sequence for every read. The output of this step is a first version of the lookup table. Errors such as barcodes assigned to multiple mutants in this raw version of the LUT are then purged in a python script collapsing all the reads, and filtering them for their confidence. Barcodes with the highest number of reads linking the barcode to the mutation are kept and if two barcodes were read the same number of times, the one with the higher quality was kept in the LUT. Then, the LUT was filtered eliminating barcodes that have the wrong length and barcodes that were read less than twice.

### Expression and labeling of myoglobin barcoded library

Aliquoted glycerol stocks of the library were thawed on ice and resuspended in liquid -TRP medium supplemented with 2% (wt/vol) glucose and Ampicillin 100 μg/ml at a starting OD of 0.1 in 50 mL. After preculture growth for 30 hours with shaking (180 rpm), appropriate volume of cells were spun down and resuspended in 100 mL at an OD 0.4 in -TRP liquid induction medium with 1.8% (wt/vol) galactose, 0.2 % glucose and Ampicillin, buffered with 100 mM sodium phosphate buffer pH 7. Cells were grown at 20°C for 48 hours with shaking at 180 rpm. Then, the culture was spun down, washed once, and resuspended in PBS BSA 0.1% (wt/vol). For cell staining, 4 *10^6^ cells were collected in a round bottom 96 well plate and incubated in 100 ul of 1:500 dilution of primary Anti Histag Antibody (stock concentration of 1.2 mg mL^−1^). Afterwards, cells were washed again with 200 ul of PBS BSA 0.1% (wt/vol) and incubated for 20 min in ice cold dilution (1:500, stock concentration 2 mg mL^−1^) of secondary goat anti-mouse antibody conjugated with Alexa Fluor 488. Then, cells were washed again twice with cold PBS BSA 0.1%.

### Tyramide labeling

For the high throughput screening of the variant and wild type activity, the protein displaying cells were subjected to a single-cell tyramide fluorophore proximity labeling technique (EP-Seq)^26^. The cells, once expressing the library, were stained by primary and secondary antibody labeling, targeting the polyhistdine tag. Then, 3 million cells were added (3000 cells/ul) to 1 mL of 50 mM sodium phosphate buffer, pH 6.1, containing 1:333 dilution of Alexa fluor^TM^ 594 Tyramid reagent (Thermo Fisher, B40957), 100 mM glucose and 62.5 nM Glucose Oxidase (from *Aspergillus niger,* Thermo fisher). The substrate concentration was chosen to yield enough signal to discriminate active from inactive variants, without saturating the yeast cell surface for the WT, with the idea that higher active variants will result in higher fluorescence after labelling reaction (see **Figure S1**). Tyramide AF594 is a fluorescently labeled phenolic substrate consisting of a tyramide moiety conjugated to an Alexa Fluor 594 dye. Upon peroxidase-catalyzed oxidation in the presence of hydrogen peroxide, the phenolic group forms a short-lived radical that covalently reacts with nearby electron-rich residues or surfaces. The exact concentration of the tyramide dye is not reported by the manufacturer. We estimated the concentration by measuring UV-VIS spectrophotometry using an extinction coefficient of ε_TyrAF594_ = 92’000. Therefore the 1:100 diluted stock solution corresponded to 3.935 uM (see **Figure S8**).

The cells were kept in suspension by shaking at 1000 rpm on a Thermomixer, while incubating for 60 min at 37°C. Then, the reaction was stopped by addition of ice cold 400 uL of PBS BSA (0.1%) supplemented with 0.05% Tween20 and subsequently washed twice by spinning cells to pellets at 13’000 g for 1 min and resuspending in 500 uL of the same PBS buffer. Then, cells were taken to fluorescent activated cell sorting (FACS) (**Figure S1**).

### Fluorescent activated cell sorting for deep mutational scanning

The capped, expression-stained and tyramide labeled library was sorted on a BD FACS Aria. The library was loaded on the cell sorter and gates were applied as follows. Gate 1 (Bin (-)) included all non expressing and non-redshifting (inactive) cells. The rest of the cells were then projected on a histogram plot displaying the ratio of the signal of the yellow over blue channel, proxying the expression normalized activity shift. On this histogram we then added three gates to capture the remaining cells by equiproportionate binning (Bin 2(+), Bin 3(++) and Bin 4 (+++) (**Figure S4**) . The same library was grown twice in biological replicates and the sorting experiment was conducted with each of the libraries, sorting each time at least 2.5 Million cells, hence covering the library 25 times. Additionally, we sorted 1000 cells from a separately grown expression culture of wild type with a known barcode in every sort tube. This internal control allows for monitoring any bias in growth conditions in between tubes. The gated populations were sorted in -TRP media supplemented with 1 % BSA to facilitate pelleting of the cells. Cells were spun down and resuspended in 10 mL -TRP medium with 2% glucose and ampicillin. The cultures were let recover and grow for 48 hours at 30°C with shaking at 180 rpm. Then, aliquots of 5 *10^7 cells were stored as glycerol stocks at -80°C.

### Illumina barcode sequencing

In order to prepare the sorted bins for high throughput sequencing, glycerol stocks (2 per bin) from both biological replicates were thawn for 5 minutes on ice and spun down at 13000 g for 1 min in a benchtop centrifuge. Then, the supernatant was removed and 250 ul of resuspension solution from bacterial miniprep kit (GeneJET Plasmid Miniprep Kit, Thermo Fisher) were added for resuspension of the pellet. Per tube, 4 ul of Zymolyase was added, and let incubate for two hours at 37 °C with shaking at 1000 rpm. After visual inspection for successful lysis of yeast cell walls (solution appears clear), 250 ul of lysis buffer and subsequently 350 ul of neutralization buffer was added, following the standard protocol for Plasmid miniprep. For better coverage of binned barcodes, two Zymoprep tubes per bin were combined in one miniprep column. After conducting the washing steps, final elution was done in 15 ul of nuclease free water. Those 15 ul then served as template for the PCR reaction, amplifying the 15 N barcode containing region in a standard 50 ul PCR reaction (NEBNext Ultra II Q5 Master Mix, New England Biolabs). 25 ul of Master Mix were added to 15 ul of template DNA with 5 ul of each primer (final conc 1 uM). For forward and reverse primers used to amplify individual binned genetic information, we used staggered primers (Ilumina_for/rev 1-4, **Table S1**), in order to provide different starting nucleotides during illumina sequencing in an else very similar amplicon. Primers were designed for use with the Illumina Nextera indexing library prep.

The amplicons for all bins were then loaded on a standard 1 % (wt/vol) agarose gel to verify correct size and then purified form cut gel using a GeneJET Gel Extraction Kit (Thermo Fisher). After cleaning and concentrating the DNA further using a DNA Clean and Concentrator-5 kit (Zymoresearch), we determined DNA concentration. In a second PCR (**Table S4**) using 1 ng of template DNA, the Nextera indexing sequence was added for all of the bins. Samples were purified with AMPure XP beads, pooled and sequenced through Novaseq 6000 Illumina sequencing.

A computational pipeline was established to process the obtained Illumina reads in order to receive the information about abundance of corresponding mutations in each bin. By using the BBMap algorithm and providing the illumina reference file containing a 15 N barcode, the sequences were mapped and alignment information stored in a sam file ^42^. Barcodes were then extracted with the pib.c script and linked through their corresponding mutation by running the rib.c script which searches for the barcodes and assigns tags according to the description in **Table S5**. Finally, all barcodes present in the LUT were sorted and grouped by identity, revealing the total number of each variant in each bin.

In a jupyter notebook (ill_tag1_bins), the grouped barcodes from all the bins were merged into one file, which was then used as the input file of the further analysis in the python script.

### Fitness score quantification

The files containing the abundance info for all barcodes in all bins was loaded in a jupyter notebook, processing the data for the generation of a fitness score for all mutants. Raw read numbers were transformed into numbers of sorted cells (*Cv*) by applying following equation, where (*Ctot*) is the total number of sorted cells per bin, (*Rv*) is the number of reads per variant and (*Rtot*) is the total number of reads per bin.

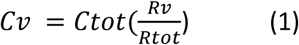

Then, for every barcode, a fitness value was determined by calculating the expected value of the fluorescent intensity across all gated bins. The total cell number for all barcodes was multiplied by the weights given through the MFI in every bin, and hence a weighted mean was determined.

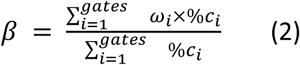

Finally, the stability fitness score was defined as the log2 of the ratio of weighted mean per variant over weighted mean of the wild type.

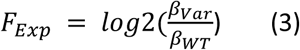

For the biochemical analysis of the library (**Figure 2**), scores from barcodes coding for the same mutation were averaged and we only considered mutations that were covered by at least 50 cells among all bins.

**Figure 2).**
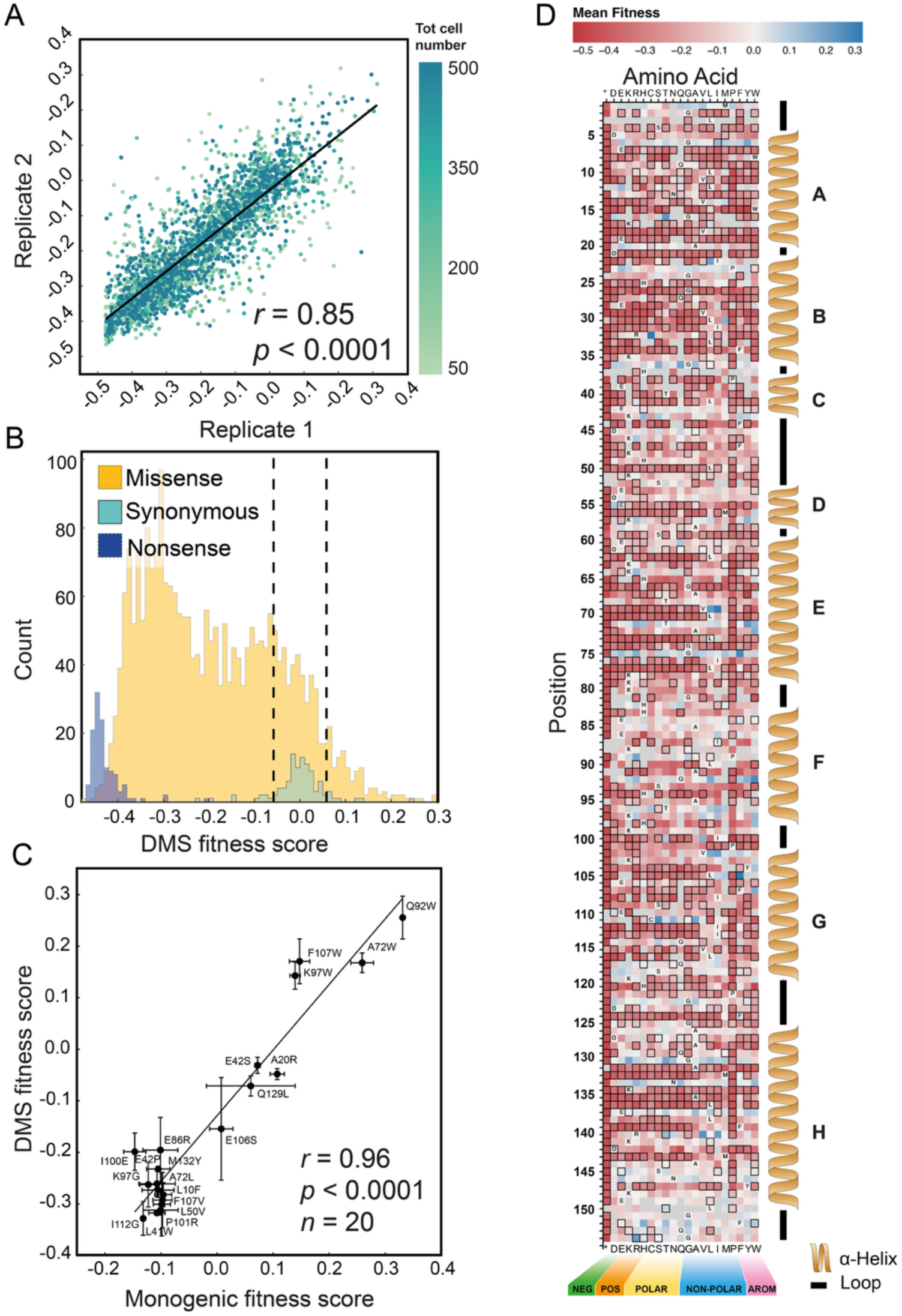
Deep mutational scanning of myoglobin peroxidase activity. **A)** Correlation of activity scores between two biological replicates. Color shading indicates total cell count per variant as indicated in the color bar. **B)** Distribution of activity scores grouped by mutation type. Missense mutations are shown in yellow, synonymous mutations in teal, and nonsense mutations in blue. Dashed lines indicate ±1 standard deviation from the mean of the synonymous codon scores. **C)** Validation of the DMS fitness scores using individually expressed monogenic variants. **D)** Heatmap of expression-normalized peroxidase activity scores for all single mutants. Amino acids are grouped by chemical class. Black borders around the squares indicate variants previously shown to have reduced stability based on a separate expression DMS screen Adapted with permission from ref [^25^]. Copyright [2026][John Wiley and Sons].

### Monogenic validation of fitness scores

In order to determine the activity score for monogenetic cultures of the variants, yeast competent cells were transformed with the respective plasmids and variants expressed on the surface as described above. Then, cells were stained with the same antibodies and protocol as used in the DMS experiment. The variants were then, along with WT, used in Tyramide labeling reactions as described above. After washing, cells were loaded on the flow cytometer and 10’000 events recorded. Since the DMS sorting was conducted in a ratio based manner in order to normalize for expression level, we used a similar scoring strategy. We set 4 gates capturing all singlet cells, as it was done in the *en masse* experiments, capturing all non expressions and nonactive cells in one gate and dividing the remaining cells in three populations. The gates were set on the wild type population to closely match the percentages obtained for wild type sequences in the DMS experiment and left constant for the whole experiment (See **Figure S4**). For each variant, the percentage of cells falling in each gate was multiplied with the mean MFI for the same gate over all variants. We determined a weighted mean for every variant by using the percentage of cells falling in each gate as weight, similarly to the procedure used in generating the high-throughput DMS score (Equation 2, where MFI corresponds to the mean of the per gate overall variants). To set the calculated score in relation with the wild type, we report the final score as the log2 values of the ratio (Equation 3).

### Pathways analysis with eMap and EHpath

We used the eMap server to find potential electron hopping pathways. For that we loaded the Alpha fold structures and used default settings for distance options (Center of mass), Surface definition (Residue depth) and amino acids taken into account (Tyrosine & Tryptophan)^43^.

For further analysis of such antioxidant rescue pathways, we used the “EHPath” python script, estimating mean residence time for available pathways, which is defined as the inverse of the transfer rate. Hence do low numbers describe fast pathways, whereas bigger values amount to faster hopping rates.

The script is fed with coordinates for an electron donor (here the heme), an electron acceptor (defined surface accessible residue) and bridging amino acid residues which are extracted by the following python command, as described in the reference^44^.

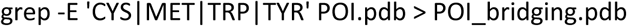

The value for the requested cut off number of maximum nodes was set to 4, indicating that each pathway consists of maximal 5 nodes. We extracted residence times for the 5 quickest pathways and set the alpha value to 1, whereas alpha is defined as

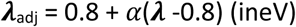

yielding ***λ***_adj_ = ***λ***, with ***λ*** being the reorganization energy and ***λ***_adj_ the adjusted reorganization energy.

### Substrate docking with Swissdock Autodock VINA

The substrate docking experiments were conducted using the Swissdock Autodock VINA engines (https://www.swissdock.ch/) ^45,46^. We submitted the tyramide fluor AF594 substrate as SMILES string: CN1C2=C(C=C3C(OC4=CC5=[N+](C)C(C)(C)C=C(CS(O)(=O)=O)C5=CC4=C3C3=C(C=CC(=C3)C(=O)NCCC3=CC=C(O)C=C3)C([O-])=O)=C2)C(CS(O)(=O)=O)=CC1(C)C and uploaded the pdb files.

We set the search box around the heme access site and set sampling exhaustivity to 10. The best ranked pose was extracted along with the calculated affinity.

For the docking studies with attracting cavities, also provided by Swisdock, we set the search space around the whole protein as a 70*70*70 Angstrom cube and set the RIC (Random initial conditions) to 5.

### Kinetics Michaelis menten analysis

Catalytic constants for RB19 and Guaiacol were determined using a TECAN UV-VIS spectrophotometer plate reader. Reaction mixtures were set up in 96 well plates and contained 1.3 uM Mb, 1 mM H_2_O_2_ and the reducing substrate in a 50 mM sodium phosphate buffer (pH 6.1). Rb19 was titrated from 5-400 uM and Guaiacol from 0.01-6 mM. Then the reactions were tracked spectrophotometrically (Decrease in absorbance at 595 nm for RB19 or increase at 470 nm for Guaiacol. The slopes were transformed in substrate turnover by means of calibration curves and the initial velocities extracted. All reactions were completed in triplicates. The reaction velocities were fitted with a Michaelis-Menten curve in Graphpad Prism.

### Michaelis-Menten analysis with tyramide decoupled assay

In order to measure michaelis menten kinetics for the actual screening substrate tyramide AF594, we design an experiment assaying the endpoint fluorescence of uninduced yeast cells containing an empty cassette. For that we prepared 96 deep well plates containing 0.27 uM (final concentration) purified soluble myoglobin in 50 mM sodium phosphate buffer (pH 6.1). Then we added the reaction mix containing the cells (20 mio/mL), 1 mM (final) H_2_O_2_ and the tyramide substrate titrated from 1 uL to 35 uL /ul. These volumina were taken from a stock of 1:100 diluted tyramide stock. The acquired stock contains 150 “slides” dissolved in 150 ul DMSO. The exact concentration cannot be stated since it is unknown. While in the DMS experiment and the decoupled cell labelling validation in **Figure 4C** the hydrogen peroxide was administered by GOx/glucose system to minimize oxidative stress for yeast cell survival, here we directly added it with the intention to lower complexity for the time sensitive experiment.

The 96 well plate was kept at 37°C with shaking at 850 rpm. Then, reactions were stopped by diluting the reaction mix adding 400 uL of PBS with 0.1 % BSA and 0.05% Tween20 supplemented with 10 mM ascorbate. The ascorbate acts as a highly concentrated reducing agent and together with the dilution effect stops the reactions reliably, as has been assayed in separate experiments. Time points were 1, 3, 5 and 10 minutes. Then the cells were washed with 400 uL of the same PBS buffer and finally resuspended in 200 uL. Then the cell’s fluorescence was assayed by means of an Attune NxT flow cytometer equipped with a plate autosampler. The median fluorescence intensity of singlet cell population was exported and used to fit the reaction speed for every concentration. Linear regression analysis was performed using GraphPad Prism software.

### Machine learning models of variant activity

#### Dataset and Preprocessing

The initial library contained 6,115 variants with up to five mutations relative to the wild-type sequence. Variants with four or five mutations were excluded from further analysis. For the remaining variants, fitness scores for expression-normalized activity were obtained from two biological replicates. A filtering step was applied, removing variants that showed an absolute difference greater than 0.1 between their replicate fitness scores to ensure data robustness. The final fitness score for each variant was calculated as the arithmetic mean of its two replicate scores. This preprocessing resulted in a dataset of 4,769 unique variants, which were used for model development.

#### Sequence embeddings

Protein sequence embeddings were generated using two pre-trained models: ESM3 “ESM3_OPEN_SMALL (esm3-open-2024-03)” and ProtTrans “prot_t5_xl_half_uniref50-enc”. For ESM3, each sequence was converted into a 1536-dimensional embedding vector, and for ProtTrans T5-XL, into a 1024-dimensional vector. In both cases, embeddings were obtained by taking the mean of the final hidden layer representations across all amino acid positions.

#### Model architectures and baselines

Several machine learning approaches were benchmarked for predicting variant fitness from sequence representations. These included Random Forest Regressor (RF) and Support Vector Regressor (SVR) using both one-hot encoded sequences and pre-trained protein language model embeddings (ProtTrans and ESM3). A Convolutional Neural Network (CNN) was also evaluated with one-hot encoded sequences (Figure 3C, left). The Multi-Layer Perceptron (MLP) architecture, when paired with ProtTrans embeddings or ESM3 embeddings, yielded the highest initial performance and was selected for further development. After comparing the performance of both embeddings on unseen variants (Figure 3C, right and Supplementary Figure S6), we selected the ESM3 model for the screening of unseen variants. The final MLP architecture consisted of an input layer with 1536 units for ESM3, followed by three hidden layers with 512, 256, and 128 units, respectively. Rectified Linear Unit (ReLU) was used as the activation function after each hidden layer. To prevent overfitting and stabilize training, Batch Normalization was applied after each hidden linear layer, and dropout with a probability of 0.6 was applied after each activation.

**Figure 3).**
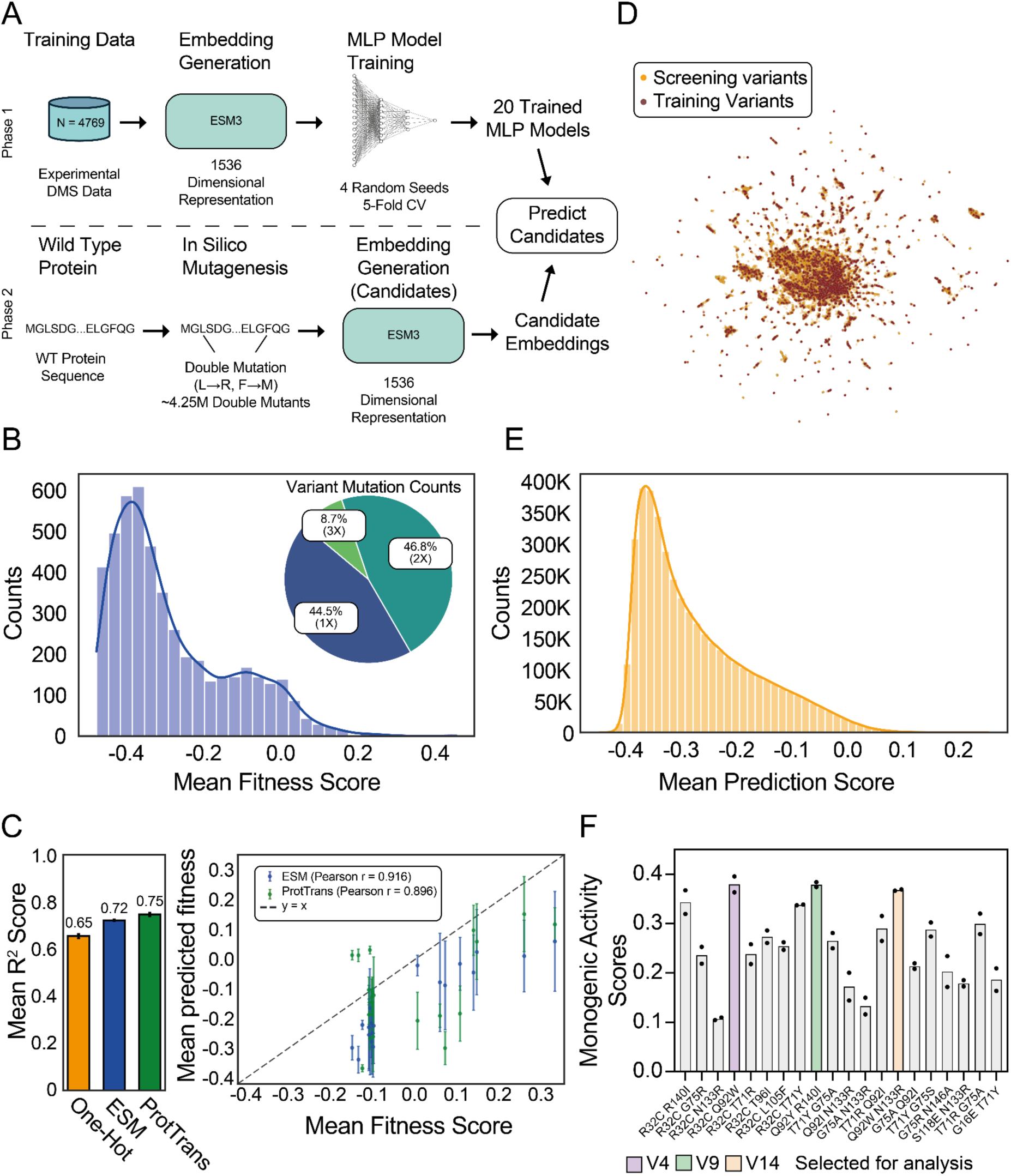
Machine learning workflow, model performance, and experimental validation of predicted high-activity double mutants. **(A)** Overview of our sequence-to-function prediction pipeline. Phase 1: Activity DMS data were used to train a feedforward neural network (multilayer perceptron) regressor using ESM3 embeddings as input features. Phase 2: An *in silico* library of ∼4.25 million double mutants was embedded using ESM3 and scored using an ensemble of 20 models. **(B)** Characteristics of the training data (N = 4,769 unique variants) after filtering out low confidence variants and higher order mutants. The histogram shows the distribution of experimental DMS scores. The pie chart indicates the proportion of single, double, and triple mutants in the training data. **(C)** Evaluation of model performance. Left: predictive performance on a held-out test set of two MLP models trained on ESM3 and ProtTrans embeddings, and a 1D convolutional neural network model trained on amino acid one-hot encodings. Bars are mean R² scores between predicted and ground truth fitness scores, computed across predictions from models trained in 5-fold cross-validation; error bars denote one standard deviation across folds. Right: Comparison of model predictions and experimentally measured monogenic activity scores for 20 variants not used in training (**Table S6**). **(D)** Two-dimensional UMAP projection of ESM3 embeddings of training myoglobin variants and the double mutants screened with the pretrained model. A random sample of 50,000 variants from the full screening library is shown in orange, highlighting extensive overlap in coverage between both libraries. **(E)** Distribution of consensus predicted fitness for the ∼4.25 million double mutants. The histogram shows the mean predicted fitness scores across all 20 trained ESM3 MLP model instances. **(F)** Experimental validation of 20 high-confidence candidate variants selected from the top 0.2% of predicted variants. All tested variants showed activity above wild type.

**Figure 4).**
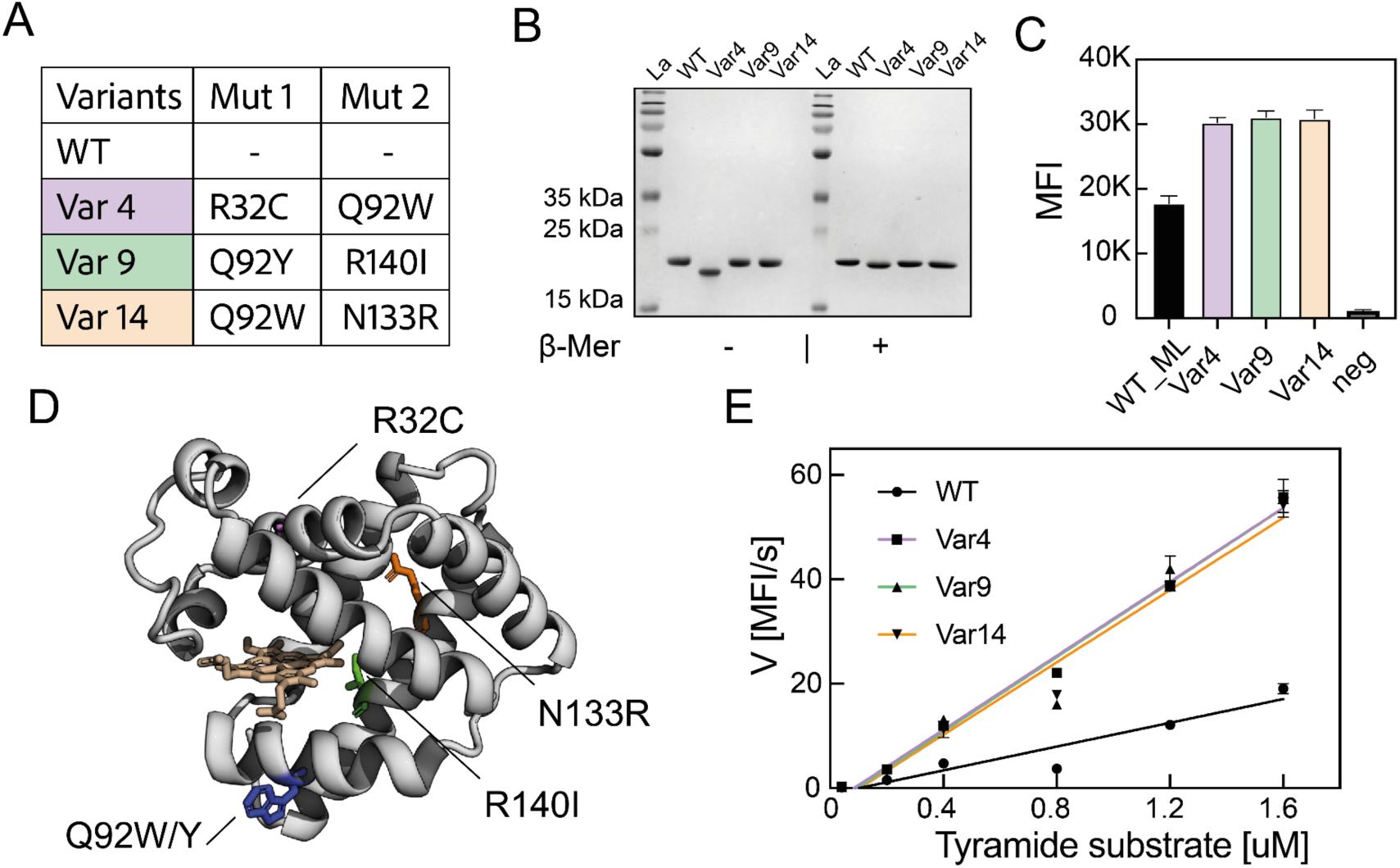
Soluble validation of ML predicted variants. **A)** Selected variants and featured mutations. **B)** 15% SDS-PAGE gel of purified soluble variants. Samples were boiled under non reducing (-) and reducing (+) conditions. **C)** Endpoint MFI of uninduced cells stained by purified double mutants with tyramide AF594. Error bars correspond to STD of triplicates and negative control contains no myoglobin in reaction mix. **D)** Structure showing positions of double mutants, according to color code introduced before. Position Q92 which is mutated in multiple variants is shown here only as Trp and in blue. **E)** Michaelis Menten analysis of soluble enzymes with tyramide. Uninduced yeast cells carrying an empty cassette were stained at different substrate concentrations and endpoint fluorescence assayed at different time points in order to get reaction velocities for variants along with WT.

#### Model training and evaluation

Models were trained using the Pytorch library. The dataset of 4,769 variants was initially divided into a training set (80%) and a fixed held-out test set (20%) using a stratified continuous split based on the distribution of mean fitness scores. A 5-fold stratified cross-validation (CV) procedure was employed on the 80% training data to train and evaluate the ESM3 MLP model. To assess model robustness and account for variations due to random weight initialization and data shuffling, the entire script, including data splitting and 5-fold CV training, was executed four times, each with a distinct random seed. This resulted in a total of 20 trained MLP models (4 seeds × 5 folds).

For each of the 20 training instances, the MLP model was trained for 500 epochs using the AdamW optimizer (learning rate = 0.0001, L2 weight decay = 1e-5) with Mean Squared Error (MSE) as the loss function. A batch size of 32 was used, and gradient norms were clipped at a maximum value of 1.0. Hyperparameters such as learning rate, optimizer choice (AdamW vs. Adam, RAdam), dropout probability (0.2-0.7), activation functions (ReLU, LeakyReLU, SiLU), and the inclusion of gradient clipping were determined through preliminary experiments using ProtTrans embeddings before finalizing the ESM3 MLP training configuration. The final optimized hyperparameters and model configurations are listed in Supplementary Table S9. During each epoch of a CV fold, model performance on the corresponding validation set was assessed using the coefficient of determination (R²) between ground truth and predicted fitness. The model state (weights) from the epoch that yielded the highest validation R² score was saved as the best model for that specific fold and seed. This best model was subsequently evaluated on the fixed 20% test set (Figure 3, left).

#### Prediction on Double Mutant Library and Candidate Selection

An in silico library of all possible double amino acid substitutions was generated from the wild-type protein sequence. After removing variants already present in the training dataset, this library contained about 4.25 million unique double mutant sequences. ESM3 embeddings were generated for this entire library. Each of the 20 trained MLP models (from the 4 seeds × 5 folds) was then utilized to predict the fitness scores for all ∼4.25 million double mutants.

To identify a set of variants with robust fitness predictions for experimental testing, we selected the top 10,000 sequences predicted by each model and identified 65 consensus sequences shared by all of the 20 prediction sets, which we deemed to be variants with high-confidence fitness predictions. From this set, 20 variants were selected for experimental validation (Figure 3F). The top four ranked variants were selected directly, and the remaining variants were chosen to maximize diversity in both, amino acid positions and mutation types. Because many of the proposed double mutants shared identical or highly similar single mutations (Table S7), we prioritized sampling a broad range of mutational combinations. The selected variants are indicated in green in Table S7.

## Results

### System setup, library construction and barcoding

We first optimized the functional display of WT hMb fused to the C-terminus of the Aga2 yeast anchor protein. We successfully detected protein display by staining the His_6_-tag at the C-terminus with a primary and secondary Alexa Fluor 488-conjugated fluorescent antibody by flow cytometry^25^. We next validated selective tyramide-based cell labeling for expressed, catalytically active variants, using negative controls either omitting H_2_O_2_ or using cells transformed with empty cassette plasmids (**Figure S1**). To generate the site saturated library of the coding region of hMb, we employed primers with nested NNK codons and barcoded each variant with a unique 15 nucleotide barcode (see Methods)^40^.

After sequencing the sorted bins, we mapped the reads via a look up table to their corresponding variants, and converted the distribution of the variants among the bins to activity fitness scores. Applying confidence filters to remove variants with low sorting or sequencing coverage, we ended up with a dataset consisting of expression-normalized activity scores for 6,115 variants bearing single and multiple mutations.

### Deep mutational scanning elucidates stability-activity trade offs

The results of the deep mutational scanning (DMS) experiment to quantify peroxidase activity were processed by computational filtering to consider only the single-site mutants (**Figure 2**). To assess reproducibility, we calculated activity scores for two biological replicates of the library, and observed a Pearson’s *r* of 0.85 (*n* = 2,661; *p* < 0.0001), indicating strong agreement between replicates (**Figure 2A**). Data points with higher cell coverage are shown in darker shades, and reflect the improved correlation for variants observed more frequently in the dataset.

We categorized the variants based on mutation type, and visualized the fitness scores in a histogram (**Figure 2B**). By definition, the WT sequence is assigned a score of 0. Synonymous mutations encoding the same amino acid sequence cluster at 0.00 ± 0.06 (n = 102, 3.83% of the single mutants). Nonsense mutations encoding stop codons (n = 123, 4.62%) exhibit the lowest activity scores of -0.43 ± 0.03, consistent with truncation of the protein chain. The largest and most informative group consists of the missense mutations (n =2,436; 91.54%), describing all single mutant variants with activity scores ranging from -0.47 to 0.3. As observed in other mutational scans, most amino acid substitutions are deleterious^26,47,48^. In our dataset, ∼88% of missense mutations reduce peroxidase activity. This fraction is higher than the ∼67% of missense mutations that decrease myoglobin expression levels^25^. This higher sensitivity to mutation likely indicates that enzymatic activity imposes stricter constraints than simple folding, making the peroxidase phenotype more vulnerable to mutational disruption than folding stability alone.

To validate the results of the DMS experiment, we conducted control assays for individual variants. We selected random sequences as well as variants with high activity scores in order to cover the full range of activity. Each variant was expressed individually on the yeast surface, and peroxidase activity was measured using the same tyramide proximity labeling protocol that was applied in the pooled screen. As in the *en masse* experiment, we calculated activity scores by quantifying expression-normalized fluorescence shifts (see Methods). The resulting values showed strong agreement with the DMS dataset (*r* = 0.96, *n* = 20, *p* < 0.0001; **Figure 2C**).

**Figure 2D** presents a heatmap of activity scores for all single-point mutants, where activity is normalized by expression level. Variants with reduced folding stability compared to wild type are shown with a black border. We estimated this stability threshold based on the distribution of synonymous mutants and defined destabilizing variants as those with expression scores < -0.06 (see supplementary info and ref ^25^). Variants without a border are either stably expressed or were not present in the expression dataset.

In addition to the destabilizing mutations discussed earlier (**Figure 2D**, shown in red with black borders), we also identified variants that are stably expressed on the yeast cell surface but lose their catalytic activity. Examples include mutations at critical positions such as the proximal and distal heme-coordinating histidines (His^94^ and His^65^, respectively). Other functionally sensitive sites include residues within helix F, a region known to support heme binding but is not required for correct folding of the apoprotein. Helix F is structurally flexible and disrupted in apo-myoglobin^49^, which makes it more tolerant to mutation compared to other helices. However, our data show that mutations in this region can severely impair peroxidase activity. This points to an activity-stability tradeoff, and underscores the importance of heme positioning and axial coordination in maintaining catalytic function, even in mutants that fold and express efficiently.

### Identification of novel and highly active variants with machine learning

We next sought to discover myoglobin variants with improved peroxidase activity by combining deep mutational data with machine learning. Using the activity DMS dataset, we trained a supervised regression model to predict peroxidase activity from protein sequences encoded using pre-trained protein language models (**Figure 3A**). We evaluated both ESM and ProtTrans embeddings given their strong performance across a variety of protein prediction tasks^50,51^. The supervised regression layer was based on a deep feedforward neural network trained on the DMS activity scores using the protein embeddings as input features. To increase the confidence of model predictions, the training data excluded variants with large variation in DMS scores across replicates (**Figure 2A** and Methods). We also excluded higher-order mutants (4x, 5x) that were poorly covered in the original DMS screen. This filtering step led to a training set with N=4,769 myoglobin variants (**Figure 3B**). After hyperparameter optimization (Supplementary Table S9), both ESM and ProtTrans models performed well on held-out test data, and both outperformed models based on one-hot amino acid encoding (**Figure 3C, left)**. We next queried both models with a set of N=20 variants for which we had independently measured monogenic activity scores (**Table S6**; Methods). Comparison of predicted and measured values indicated that ESM embeddings provided better out-of-distribution accuracy. Based on this result, we selected the ESM embeddings for the final computational screen (**Figure 3C, right**).

Given that the training data was highly enriched in double mutants (46.8%) and included only a small fraction of triple mutants (8.7%), we restricted the prediction screen to double mutants that were not present in the training set. This increased reliability of model predictions. Moreover, a two-dimensional projection of both the training and query sequences showed good overlap, which further supports the robustness of the predictions despite the relatively limited coverage of the training data (**Figure 3D**). We embedded a total of N=4,250,505 double mutants using ESM and queried an ensemble of 20 regressors, each trained using five-fold cross-validation and four random seeds for weight initialization. The predicted activity scores for these unseen double mutants followed a distribution similar to that of the training data (**Figure 3E)**. For experimental validation, we selected 20 from a set of 65 consensus hits that scored in the top 0.2% (N=10,000 variants) across the model ensemble (**Table S7**). We note that all of the 20 double mutants for which single-mutant activity measurements were available consisted of combinations of variants with higher activity than WT. The predicted activity of double mutants showed a strong correlation with the sum of the corresponding single-mutant scores, indicating that the model largely captures additive effects and predicts only limited epistatic interactions (**Figure S10**). All 20 of these tested variants showed improved peroxidase activity over the wild type. This perfect success rate demonstrates the effectiveness of our machine learning-guided approach for identifying highly active myoglobin variants.

### Analysis of best performing ML predicted sequences as soluble enzymes

After showing that the top machine learning-predicted variants all exhibited higher peroxidase activity than WT myoglobin when displayed on the yeast surface, we further characterized the top three candidates as purified soluble enzymes. These variants, referred to as Var4, Var9, and Var14, were selected based on their top ranked monogenic activity scores and were expressed in *E. coli* along with WT myoglobin. The mutational compositions of these variants are shown in **Figure 4A**, and their positions are mapped onto the 3D structure in **Figure 4D**. Mutations were modeled using AlphaFold and visualized in PyMol as sticks. We obtained the purified protein and could verify by means of reducing and non-reducing SDS page that the R32C mutation, known from our prior study^25^, forms a disulfide bond in the ML-predicted double mutant (**Figure 4B)**. The non-reduced form of the protein migrates faster due to the intramolecular disulfide with C111, while addition of β-mercaptoethanol eliminates this change in electrophoretic mobility.

To test peroxidase activity for the soluble variants, we adapted the tyramide labeling assay to a soluble format. Employing the same reaction mixture as with yeast displaying proteins, we stained uninduced yeast cells carrying an empty cassette plasmid by administering soluble hMb variant enzymes. The endpoint fluorescence data (**Figure 4C**) confirmed that all three double mutants produced higher signal than wild type, reproducing the high activity ranking observed in the yeast-displayed format. Negative controls lacking any enzyme showed minimal background fluorescence.

To gain a deeper understanding of the kinetics of these improved variants, we performed a Michaelis-Menten analysis using the same endpoint-based tyramide labeling assay. Reaction velocities were measured at varying substrate concentrations by stopping the reaction at different timepoints and fitting linear regressions to the endpoint mean fluorescence intensities (MFIs) (see methods). **Figure 4E** shows the resulting velocity curves for WT and the three selected double mutants. Due to the high cost of tyramide, we were unable to reach substrate saturation. Therefore, a linear approximation of the Michaelis-Menten model was used in the low-substrate regime to estimate catalytic efficiency. All three double mutants showed slopes more than three times steeper than WT, indicating high catalytic rates at the concentrations tested, including under the standard library screening concentration which corresponds to 1.2 uM.

In all three of these improved variants (Var4, Var9, and Var14), residue Q92 is mutated to either tyrosine or tryptophan, suggesting that the introduction of a redox-active residue at this surface site contributes to enhanced activity. The R32C mutation found in Var4 likely serves a stabilizing role that supports acquisition of secondary activity-enhancing mutations. This stabilizing effect of R32C was presumably the reason it was frequently found among top ML-ranked double variants. To further explore the catalytic behavior of the selected variants, we expanded our kinetic analysis to include alternative peroxidase substrates, specifically reactive blue 19 and guaiacol. The results are presented in **Figure S2** and discussed in the Supplementary Note **SN1**.

### Strategically placed Trp wires boost activity towards bulky substrates

Among the machine learning-predicted variants discussed above, we observed a clear enrichment of mutations that introduce tyrosine or tryptophan. Beyond the frequently occurring R32C disulfide variant^25^ found in many of the top double mutants (**Table S7**), four of the five most active candidates contain either a tyrosine at position 71 or a tyrosine or tryptophan at position 92. This trend is also observable across the whole DMS dataset, where single-point variants containing these substitutions often show higher peroxidase activity. This can be seen in the heatmap in the two rightmost columns in **Figure 2D**. Conversely, mutations that remove native tyrosine residues tend to show very deleterious effects on activity (**Figure S3**). For the two native tryptophan residues we cannot estimate their role in activity since both are elementary for stability and do not tolerate substitution.

When comparing the effects of mutations on protein stability and catalytic activity, we find that aromatic amino acids are particularly important for maintaining function. In many cases, mutations that are tolerated in the expression screen exhibit reduced activity when an aromatic side chain is removed. Wild type tyrosine, histidine and phenylalanine residues tolerate some substitutions, but these almost always come at the expense of catalytic efficiency.

These findings support a broader role for redox-active aromatic residues in modulating myoglobin peroxidase activity. It has been reported that many oxidoreductases possess clusters or chains of tyrosine and/or tryptophan residues that serve as hole hopping relay stations, providing alternative electron transfer pathways to mitigate oxidative stress and preserve function.^21,52^ Similarly, dye-decoloring peroxidases use surface exposed aromatic residues as stepping stones for oxidation of bulky substrates that cannot pass through the heme access tunnel.^53,54^ Such work has inspired others to use surface tryptophans and tyrosines in myoglobins to enhance dye decoloring peroxidative activity in myoglobins.^12,13^

Consistent with this rationale, our data show increased peroxidase activity for variants introducing tyrosine or tryptophan at surface-accessible positions. In many cases, these substitutions are interchangeable, supporting the idea that their redox potential rather than specific side-chain orientations are beneficial to catalysis (e.g., positions 71, 92, 137, 146 or 152). We visualized this trend in **Figure 5A** by mapping all single tryptophan substitution scores onto the protein surface. Variants that improve activity are shown in blue, and deleterious substitutions are shown in red. Positions where tryptophan residues increase the peroxidase activity are placed around the heme active site on the surface of the protein. This spatial arrangement suggests that these engineered residues facilitate electron transfer for substrates like tyramide, which are too bulky to access the heme directly. We note that the native tyrosine and tryptophan residues are not surface exposed (SASA W8: 4, W15: 2, Y103: 18, Y146:0, where 0 is buried and 100 equals fully exposed). These observations support the idea that enhanced labeling efficiency arises from introduction of redox relays close to the substrate interface.

**Figure 5).**
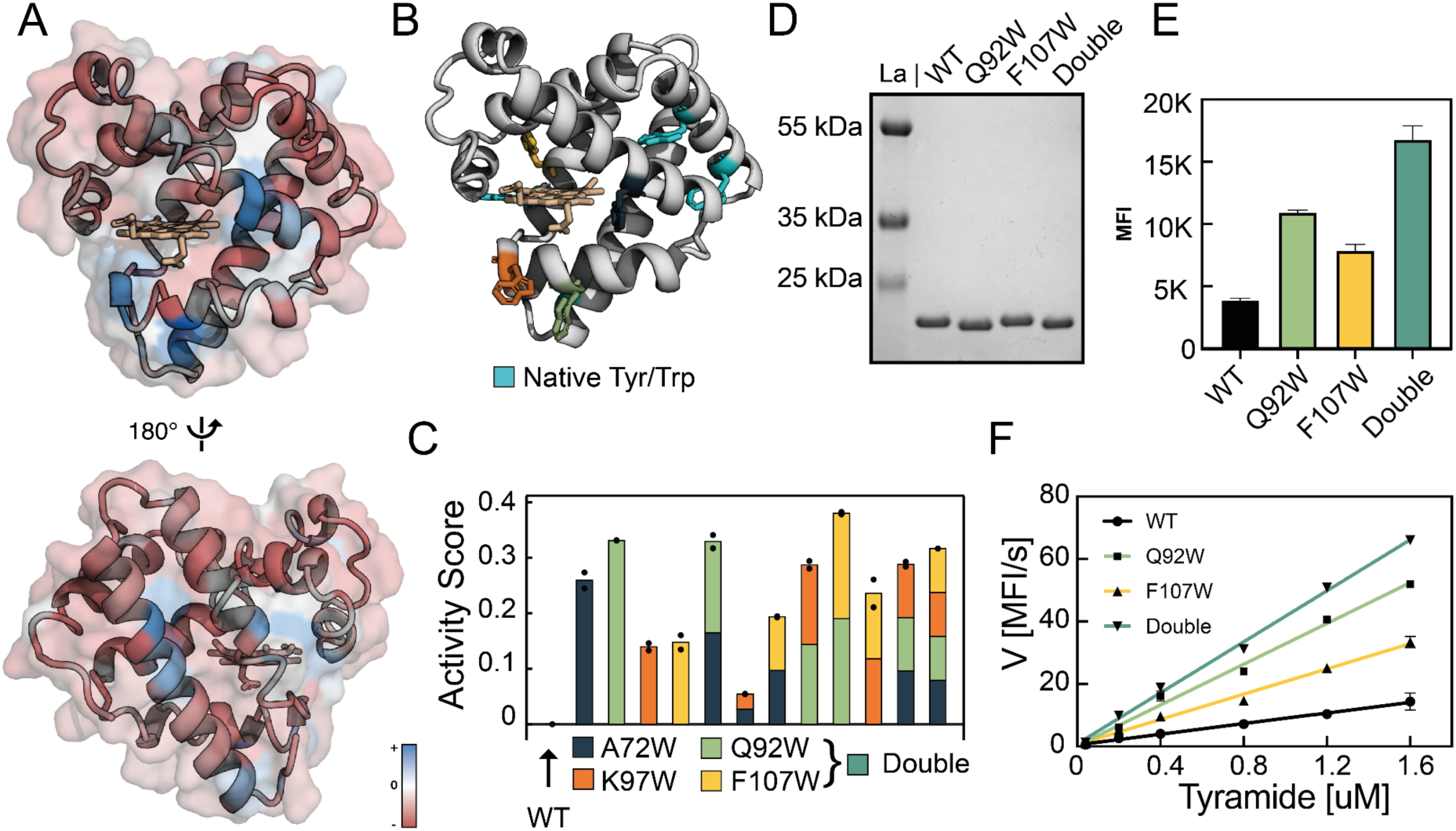
Tryptophan-mediated enhancement of peroxidase activity. **A)** Myoglobin 3D structure with residues colored by single mutant fitness scores for TRP substitutions. Positions without measurements are shown in white as cartoon loops. The heme cofactor is shown in white. **B)** Residues selected for combinatorial Trp-library are shown as sticks and colored according to the legend in panel C. **C)** Barplot of monogenic activity scores for individual TRP mutations and their combinations. **D)** SDS-PAGE gel analysis of purified protein variants. **E)** Decoupled cell labelling assay. Endpoint MFI values are shown for uninduced yeast cells containing an empty cassette plasmid, stained with soluble enzymes and the tyramide fluorophore. **F)** Reaction velocities for soluble enzyme variants measured using a time-dependent tyramide assay. Data represent the linear region used to approximate catalytic efficiency.

To examine whether the activity-enhancing effects of individual tryptophan mutations could be combined synergistically, we designed a small combinatorial library targeting the four positions with the highest fitness scores in the single-mutant DMS dataset, which were A72W, Q92W, K97W and F107W. Notably, substitution A72W, which yields one of the most active variants in the library, was the only substitution at that position 72 that retained or improved peroxidase activity, while all others were neutral or deleterious. Furthermore, since most mutations at A72 were neither strongly destabilizing nor beneficial in terms of expression stability^25^, this position seems to be functionally specific to peroxidase activity.

The selected variants were modeled using AlphaFold and visualized in PyMol^55^ (**Figure 5B**). Residues were color coded according to the legend in **Figure 5C**, where native tyrosine and tryptophan residues are shown in cyan. We synthesized these combinatorial variants using site-directed mutagenesis and tested their peroxidase activity in the yeast surface display system (**Figure 5C**) using the same WT-normalized scoring method as described previously (see **Figure S4**). Consistent with the prior DMS data and ML predictions, all single TRP variants outperformed WT myoglobin (**Figure 5C**). The combinatorial mutants also exhibited improved activity relative to WT, although the effects were more variable. While most combinations did not show improvement upon introduction of additional TRP residues, the impact of F107W was dependent on which TRP mutation it was paired with. In combination with A72W, activity declined, whereas when combined with double mutant Q92W it was amongst the very best variants found in this study.

In addition to tyrosine residues, tryptophans possess electron-rich aromatic side chains that can under certain conditions undergo oxidative modification by peroxidase-generated radicals, especially under high peroxide or high substrate conditions^56^. To ensure that the improved labeling signal observed in our TRP-substituted mutants was not due to the introduction of these additional labeling sites, we performed the decoupled labeling assay with soluble myoglobin variants as described above. We purified WT, Q92W, F107W and Q92W/F107W variants as soluble proteins (**Figure 5D)** and used them to stain uninduced yeast cells containing only an empty cassette plasmid under otherwise identical reaction conditions. This decoupled labelling assay eliminated any artefacts due to the display format, and allowed direct attribution of labeling signal to peroxidase activity **(Figure S9)**. The same performance trend (**Figure 5E**) observed in the display assay was found in this decoupled format, confirming that the enhanced signal was the result of increased enzymatic rate rather than TRP-modification on myoglobin itself.

As presented in **Figure** for the ML predicted variants, we quantified reaction velocities at different tyramide concentrations as well for the tryptophan mutants. As shown in **Figure 5F**, the double mutant Q92W/F107W exhibits a 4.9-fold increase in reaction velocity relative to the wild type at the substrate concentration used in the DMS assay. Interestingly, although the single mutants Q92W and F107W each individually enhance reaction velocity by factors of 3.9 (DMS: 0.26) and 2.4 (DMS: 0.17), respectively, their combined effect in the double mutant was not fully additive, indicating some negative epistatic effects.

We further attempted to explain observed trends for mutants by analyzing the potential electron transfer pathways employing the published tools eMap and EHPath. eMap is a python based web application that predicts possible electron or hole transfer channels from pdb files based on graph theory. Shortest path algorithms are used to estimate shortest pathways from user-specified hole donor (here heme) to the surface of the protein, assigning scores to all pathways^43^. EHPath is another python module estimating and ranking mean residence times of a transferring charge along such hopping pathways^44^. The settings used are described in the method section and the results are presented in **Figure S5** and discussed in supplementary notes **SN2**. We were interested in studying the additivity of the double mutant Q92W/F107W in more detail and hence used purified variants to test the activity in Michaelis-Menten analysis. Equally to the machine learning predicted variants above, we switched to reactive blue 19 as substrate for this experiment. The results including the fits and extracted kinetic parameters are shown in **Figure S7**. We see that the trend observed in the monogenic tyramide scores as well as the decoupled cell labelling assay holds true also for Rb19 dye, with both single mutants being more active than WT and the double mutant benefiting from additivity of composing mutants.

Next, we assayed how the mutations would act on peroxidase activity towards smaller substrates that might fit in the active site crevice. We performed a Michaelis-Menten assay using guaiacol for the selected variants along with WT myoglobin and saw that in fact the activity towards this smaller substrate is similar for all variants tested (**Figure 4E**). We used molecular docking to further validate our hypothesis, differentiating between bulkier substrates that benefit from a surface radical and smaller ones that do not. We find that for WT, Q92W and F107W the binding site for guaiacol is identical and directly adjacent to the heme cofactor **(Figure S7 F**). Contrary to the machine learning variants, here we study single mutations, allowing for direct allocation of observed effects. Considering that we found improvement towards guaiacol for the machine learning predicted variant 9 which contains the R140I and Q92Y mutations, we suspect that especially the R140I mutation is leading to enhanced guaiacol reactivity, while the tyrosine mutation, similar to the tryptophan substitutions are beneficial for bulky tyramide oxidation.

Lastly, we cross-referenced our deep mutational scanning data with variants reported in gnomAD to identify mutations in myoglobin observed in human clinical populations. We annotated these clinically observed variants with their stability and activity scores derived from our screening assays (**Table S8**). Although our peroxidase screening assay used a non-physiological substrate, these annotations may still provide useful insights into naturally occurring variants. Notably, several clinically observed variants show substitution of wild-type residue with tyrosine or tryptophan, suggesting potential impacts on enhancing oxidation of bulky substrates.

## Discussion

In this study we provide a comprehensive map of the peroxidase activity fitness landscape of human myoglobin, utilizing the high-throughput EP-Seq platform. We highlight regions of activity-stability trade-offs, and show global trends of amino acid groups as well as single mutations with enhanced activity. By leveraging the extensive labeled mutant library, we integrate high-throughput DMS with ML to successfully train a predictive model and identify novel double mutants with elevated peroxidase activity. Notably, all 20 ML-predicted sequences were substantially more active than wild type (WT).

The most promising sequences were expressed as soluble proteins to assess whether their improved activities observed on the yeast cell surface translated to the soluble format. When tested at the same substrate concentration used in the initial library screening, these variants exhibited over threefold higher catalytic efficiency compared to WT. Analysis of the machine learning dataset pointed toward the introduction of oxidizable amino acids such as tyrosine and tryptophan as a key driver of enhanced activity with the bulky tyramide substrate. We validated this hypothesis by constructing a small focused combinatorial library based on top-performing tryptophan mutations. Among these, surface-exposed residues such as Q92 and A72 substantially boosted activity, supporting the hypothesis that these residues serve as redox-active relay stations and facilitate long-range electron transfer. In particular, the combination of mutations F107W and Q92W increased tyramide oxidation by nearly fivefold in assays with soluble proteins.

These findings demonstrate that activity fitness scores derived from the EP-Seq deep mutation scanning platform reliably predict activity trends in soluble enzymes, providing a key validation of this platform. Beyond methodological relevance, the insights reported will find broad applications ranging from engineering bulky dye decoloring peroxidases for industrial use to providing knowledge about mutations that alter peroxidase activity of globins, including hemoglobin based oxygen carriers (HBOCs)^57^. Surface-exposed tyrosine residues have already been shown to accelerate reduction by physiological reductants such as ascorbate in hemoglobin^58^. The strategy developed here could guide future engineering efforts to identify mutations that tune redox activity for safe HBOCs. Finally, we establish a combined approach of high-throughput screening and machine learning to expedite enzyme engineering using yeast surface-displayed libraries, with results directly transferable to soluble enzymes.

Finally, this work underscores the utility of combining high-throughput experimental fitness landscapes with pretrained protein language models to drive hypothesis generation, prioritize variants, and ultimately expand the functional repertoire of enzymes. The ability to use ML predictions to successfully guide mutational searches across vast sequence space represents a generalizable framework for accelerating biocatalyst development.

## Supporting Information

Additional experimental and computational details, materials and methods. Supplementary figures and tables.

## Author contributions

**C.K.** and **M.A.N.** conceived the study and drafted the manuscript. **C.K.** carried out the practical work and computational analyses. **A.D.** carried out the machine learning and variant prediction. **R.V.** contributed to the conceptualization and optimization of the EP-Seq experimental and computational workflow. **D.A.O.** designed the machine learning work. **M.A.N.** secured funding and administered the project.

## Acknowledgements

This work was supported by the University of Basel, ETH Zurich, and the Swiss National Science Foundation (200021_191962). AD and DAO were supported by a UKRI Engineering Biology Mission Award CYBER under BBSRC grant BB/Y007638/1.

## TOC Graphic

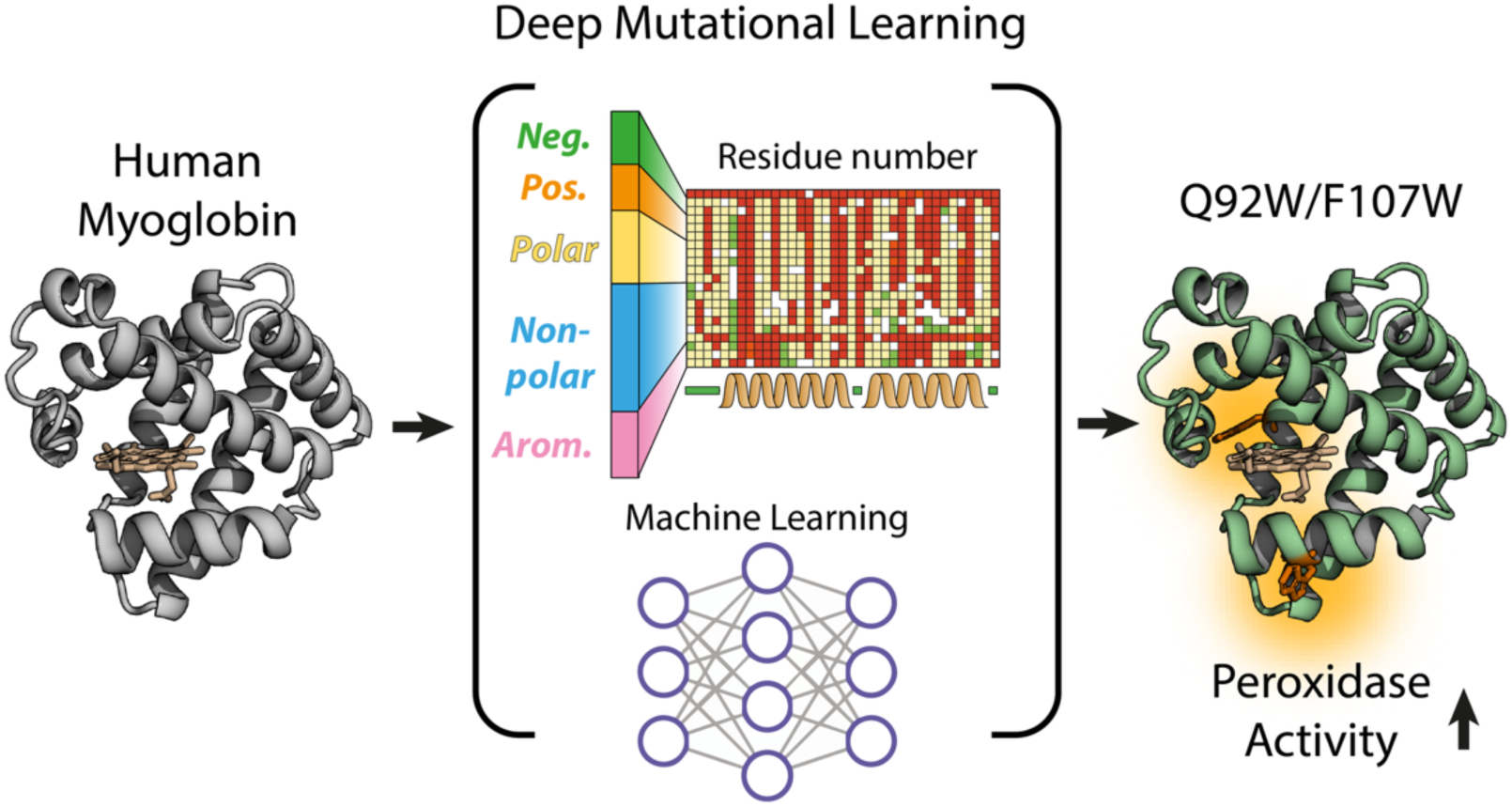

