## Supplementary Information for "Signatures of Electron-hole Hopping in Myoglobin Peroxidase Activity Revealed by Deep Mutational Learning"

### Supplementary Notes

#### SN1 - Machine learning predicted variants under substrate variation

After testing the predicted double mutants with the screening substrate, verifying their superior activity, we switched to reactive blue 19, an extensively used industrial dye, to study enzyme kinetics in more detail. Oxidation of this substrate is spectrophotometrically traceable ($\varepsilon_{595}$ = 10,000 $M^{-1}cm^{-1}$)[54] and is of biotechnological interest as dye decolorization in wastewater treatment. Further, this dye is more bulky than standard peroxidase substrates like DMP, guaiacol or ABTS hence resembling more the larger screening substrate tyramide AF 594. The Michaelis-Menten plots are presented in **Figure S2** and interestingly reveal a different scenario than for the tyramide substrate.

Variant 9 is by far the fastest enzyme ($V_{max,\ V9}$ = 115.6 ± 9.98 nM/s), exhibiting more than double the reaction velocity of WT ($V_{max,\ WT}$ = 49.93 ± 3.38 nM/s). Then, Var14 follows with $V_{max,\ V14}$ = 67.47 ± 5.26 nM/s. Striking however is that also the $K_m$ is increased to $K_{m,\ V9}$ = 244.36 ± 34.18 uM & $K_{m,\ V14}$ = 130.61 ± 15.31 uM compared to $K_{m,\ WT}$ = 82.02 ± 16.22 uM for WT, putting the catalytic efficiency into perspective. For Var4, we obtain similar kinetic constants than WT $V_{max,\ V4}$ = 50.80 ± 2.90 nM/s & $K_{m,\ 4}$ =83.71 ± 11.15 uM . This suggests while Var 4 showed enhanced activity in the original screen on yeast as well as in the decoupled tyramide assay, its improvement does not extend to other substrates like RB19, indicating substrate specific effects. Lastly, we tested the variants along with WT for oxidation of guaiacol. The results are shown in **Figure S2** and demonstrate that also for this substrate variant 9 is the most active, not only increasing Vmax but also decreasing Km towards this substrate. Together, these results suggest that for the tyramide substrate the substitution of glutamine to either tyrosine or tryptophan makes the biggest impact, resulting in comparable catalytic efficiencies for all variants. For other substrates however, the secondary mutation becomes more important, either removing the increasing effect (V4 and V14 with guaiacol and V4 with RB19) completely or attenuating its effect (V14 with RB19).

However, any comparison must be made with caution, as it involves structurally different substrates and lacks complete Michaelis-Menten fitting data for the tyramide substrate.

#### SN2 - Electron pathway and docking studies

Trying to elucidate the mechanism for improved reactivities of the found mutations, we made use of the eMap server, a publicly available algorithm (https://emap.bu.edu/single) identifying potential electron hopping pathways, based on the distance and number of aromatic stepping stones. We obtained scores ranking found pathways for all variants investigated and present them in **Figure S5 A**[43]**.** For the WT, no surface accessible tyrosine or tryptophan could be obtained, hence no pathway was suggested. We further employed EHPaths, a python script predicting mean residence times of electron hopping pathways[44] in oxidizable amino acid residues within proteins. Mean residence times of pathways located before are shown in the table in **Figure S5 A.** For the analysis, csv files of mutants were prepared as described in ref [44]. The settings used are described in the method section. The sequence of the eMap scores and the EHPath time predictions are consistent except for the F107W variant where the pathway length is ranked the longest by eMap, whereas the residence time is estimated to be shorter than for the pathway enabled by the Q92W mutation. Striking however is that the ranking from the tyramide monogenic score, shown in **Figure 4C**, is not agreeing with both hopping pathways prediction tools. This hints to the fact that not alone the charge transfer reaction but also the properties of the substrate and hence its binding to the enzyme heme or amino acids based active site, defines the activity [9].

We thus decided to employ Swissdock Autodock VINA in order to investigate substrate binding by molecular docking. In **Figure S6A** the best pose of the docking runs are presented along with their estimated affinity. In **Figure S6B** the chemical structure of tyramide AF594 is shown, which was used for docking studies. For all mutants except the A72W variant, the poses are almost identical and calculated binding energy is comparable around -8.2 kcal/mol, whereas the bulky tryptophan on helix E at position 72 hinders the substrate from taking this pose and increases binding energy to -7.4 kcal/mol. While that could explain why the supposedly faster charge transfer pathway introduced by A72W is not the most active, it does simultaneously indicate that the other mutations do not strongly affect binding around the heme site and hence other effects must lead to observed results. So does for instance show another top ranking docking pose suggest for Q92W that electron hopping could be facilitated by pi stacking of the tyramide moiety and the introduced tryptophan as shown in **Figure S5 B** (distance between planes = 3.8 Å).

#### Supplementary Tables

**Table S1)** Primers used for linkage of molecular barcodes to the myoglobin mutant variants. NNK nested primers used for the generation of the library are provided at: https://zenodo.org/uploads/16781055

| | |
|---|---|
| Barcoding Primer P1 | CGATTTTGTTACATCTACACTGTTGTTATCAGATCAGCGGGTTTAAAC |
| Barcoding Primer P2 | GCGCGGCCTTTTGCCCTGCAGGCCNNNNNNNNNNNNNNNAGGGAACAAAAGCTGG<br>CTAGTACGG |
| PacBio 1 | TTACGGTTCCTGGCGGCCGCG |
| PacBio 2 | GTCGATTTTGTTACATCTACACTGTTGTTATCAGATCAGCGGG |
| Illumina_for1 | TCGTCGGCAGCGTCAGATGTGTATAAGAGACAGGGCCTTTTGCCCTGCAGGC |
| Illumina_for2 | TCGTCGGCAGCGTCAGATGTGTATAAGAGACAGTGGCCTTTTGCCCTGCAGGC |
| Illumina_for3 | TCGTCGGCAGCGTCAGATGTGTATAAGAGACAGATGGCCTTTTGCCCTGCAGGC |
| Illumina_for4 | TCGTCGGCAGCGTCAGATGTGTATAAGAGACAGCATGGCCTTTTGCCCTGCAGGC |
| Illumina_rev1 | GTCTCGTGGGCTCGGAGATGTGTATAAGAGACAGGGAGGAGAGTCTTCCTTCGGAGG<br>G |
| Illumina_rev2 | GTCTCGTGGGCTCGGAGATGTGTATAAGAGACAGTGGAGGAGAGTCTTCCTTCGGAGG<br>G |
| Illumina_rev3 | GTCTCGTGGGCTCGGAGATGTGTATAAGAGACAGATGGAGGAGAGTCTTCCTTCGGAG<br>GG |
| Illumina_rev4 | GTCTCGTGGGCTCGGAGATGTGTATAAGAGACAGGATGGAGGAGAGTCTTCCTTCGGA<br>GGG |

**Table S2)** Computational workflow. Steps and respective commands to map the PacBio reads to the reference and extract the molecular barcode to generate the final look up table (LUT). The raw sequencing files as well as the computational scripts are provided at: https://zenodo.org/uploads/16781055.

| **Step** | **Tool** | **Command** |
|---|---|---|
| Mapping of the PacBio read to the reference file. | Minimap2<br>(version 2.19) | **minimap2 --cs -ax map-hifi** Myo_ref.fa pacbio.fastq **>** aln.sam |
| Extracting barcode sequence and quality along with resp mutations and indels. | ppba C script<br>libscodon.h library | **./ppba** Myo_ref.fa aln.sam **>** pre_lut.tsv |
| Filtering and cleaning of LUT | generate LUT.ipynb<br>Python script | |

**Table S3)** PCR conditions for barcode amplification

| Step | Temp. [°C] | Time [s] | Cycles |
|---|---|---|---|
| Initial Denaturation | 98 | 30 | 1 |
| Denaturation | 98 | 10 | 20 |
| Annealing | 72 | 30 | |
| Extension | 72 | 10 | |
| Final extension | 72 | 120 | 1 |

**Table S4)** PCR conditions for Illumina Nextera Indexing

| **Step** | **Temp. [°C]** | **Time [s]** | **Cycles** |
|---|---|---|---|
| Initial Denaturation | 98 | 30 | 1 |
| Denaturation | 98 | 30 | 8 |
| Annealing | 55 | 30 | |
| Extension | 72 | 30 | |
| Final extension | 72 | 120 | 1 |

**Table S5)** Pipeline for Illumina read processing. All bins were processed separately through the individual owing steps. The raw sequencing files as well as the computational scripts are provided at: https://zenodo.org/uploads/16781055.

| **Step** | **Tool** | **Command** |
|---|---|---|
| Sequence alignment of illumina reads to reference file. | BBMap<br>cite | bbmap.sh in=*.fq ref=reference.fa out=*.sam |
| Extracting barcode sequence and respective quality score for the illumina reads | pib.c script<br>(process illumina barcode) | pib *.sam > *.fq |
| Align the extracted barcodes to the previously processed look up table and assign tags (0 = not found, 1 = found, 2 = Read quality below Q20, 3 = Barcode length deviates from 15). | rib.c script<br>(read illumina barcode) | rib -t lut.ts -q 20 *.fq > *.tsv |
| Filtering for found variants, sort them alphabetically and group them by identity counting the number of barcodes per bin. | | grep ^1 *.tsv\|sort\|uniq -c\|sed -E 's/^ *//; s/ /\t/' > t1sct_*.tsv |

**Table S6)** 20 tested monogenic variants and their corresponding DMS activity score.

| Nr | Variant | Score |
|---|---|---|
| 1 | I100E | -0.146674 |
| 2 | I112G | -0.131686 |
| 3 | K97G | -0.122555 |
| 4 | L41W | -0.107251 |
| 5 | M132Y | -0.106929 |
| 6 | A72L | -0.105029 |
| 7 | E42P | -0.104801 |
| 8 | E86R | -0.100954 |
| 9 | P101R | -0.100502 |
| 10 | L50V | -0.098388 |
| 11 | F107V | -0.097246 |
| 12 | L10F | -0.095855 |
| 13 | E106S | 0.007579 |
| 14 | Q129L | 0.060239 |
| 15 | E42S | 0.072335 |
| 16 | A20R | 0.107461 |
| 17 | K97W | 0.139341 |
| 18 | F107W | 0.147373 |
| 19 | A72W | 0.258996 |
| 20 | Q92W | 0.331142 |

**Table S7)** Table of top scoring predicted double mutants which are found amongst top 10'000 of all prediction folds. Selected variants for monogenic testing are highlighted in green. Variants selected for purification are shown with orange numbers.

| Rank | Mut1 | Mut2 | Pred. Score | Rank | Mut1 | Mut2 | Pred. Score |
|---|---|---|---|---|---|---|---|
| 1 | R32C | R140I | 0.2510 | 34 | R32C | N133H | 0.1893 |
| 2 | R32C | G75R | 0.2313 | 35 | R32C | T96V | 0.1811 |
| 3 | R32C | N133R | 0.2292 | 36 | R32C | T96M | 0.1789 |
| 4 | R32C | Q92W | 0.2280 | 37 | Q92Y | R140I | 0.1732 |
| 5 | R32C | Q92Y | 0.2226 | 38 | T71Y | G75A | 0.1722 |
| 6 | R32C | G75S | 0.2217 | 39 | Q92I | N133R | 0.1662 |
| 7 | R32C | T71R | 0.2194 | 40 | G75T | N133R | 0.1590 |
| 8 | R32C | T96I | 0.2162 | 41 | G75S | N133R | 0.1558 |
| 9 | R32C | D45P | 0.2158 | 42 | G75A | N133R | 0.1549 |
| 10 | R32C | G75A | 0.2154 | 43 | G36L | R140I | 0.1549 |
| 11 | R32C | G36C | 0.2144 | 44 | T71R | Q92I | 0.1507 |
| 12 | R32C | G75K | 0.2136 | 45 | T71Y | G75T | 0.1503 |
| 13 | R32C | Q92I | 0.2091 | 46 | Q92W | N133R | 0.1499 |
| 14 | R32C | L105F | 0.2085 | 47 | G75R | N133I | 0.1462 |
| 15 | G6Q | R32C | 0.2084 | 48 | G75R | N133M | 0.1452 |
| 16 | R32C | G75T | 0.2079 | 49 | G75A | Q92I | 0.1431 |
| 17 | R32C | Q92R | 0.2053 | 50 | G75R | N133F | 0.1415 |
| 18 | R32C | G75L | 0.2052 | 51 | T71Y | G75S | 0.1409 |
| 19 | R32C | R140V | 0.2045 | 52 | R32A | G75R | 0.1379 |
| 20 | R32C | N133F | 0.2036 | 53 | G75R | N146A | 0.1375 |
| 21 | G6N | R32C | 0.2034 | 54 | S118E | N133R | 0.1360 |
| 22 | G6E | R32C | 0.2020 | 55 | G75N | N133I | 0.1337 |
| 23 | G6S | R32C | 0.1976 | 56 | G75A | N133M | 0.1328 |
| 24 | R32C | N133Y | 0.1972 | 57 | T71R | G75A | 0.1321 |
| 25 | G6H | R32C | 0.1959 | 58 | G75S | Q92I | 0.1310 |
| 26 | I22L | R32C | 0.1947 | 59 | G75A | N133Y | 0.1275 |
| 27 | R32C | T96L | 0.1946 | 60 | G75K | N133V | 0.1249 |
| 28 | G6M | R32C | 0.1932 | 61 | G75K | N133I | 0.1222 |
| 29 | R32C | T71Y | 0.1929 | 62 | G75S | N133Y | 0.1188 |
| 30 | R32C | N133I | 0.1908 | 63 | G75T | N133Y | 0.1174 |
| 31 | G16D | R32C | 0.1903 | 64 | G16E | N133R | 0.1134 |
| 32 | R32C | Q92F | 0.1898 | 65 | G16E | T71Y | 0.1105 |
| 33 | R32C | N146H | 0.1894 | | | | |

**Table S8)** Variants found in GNOMAD database with their annotated expression and activity scores.

| GNOMAD | Wild type | Position | Mutation | Stability Score | Activity Score | GNOMAD | Wild type | Position | Mutation | Stability Score | Activity Score |
|---|---|---|---|---|---|---|---|---|---|---|---|
| W15S | Trp | 15 | Ser | -0.465 | -0.431 | L30F | Leu | 30 | Phe | 0.095 | -0.140 |
| L136P | Leu | 136 | Pro | -0.547 | -0.427 | L138V | Leu | 138 | Val | -0.032 | -0.140 |
| G81W | Gly | 81 | Trp | -0.075 | -0.418 | K64E | Lys | 64 | Glu | -0.110 | -0.133 |
| V29G | Val | 29 | Gly | -0.492 | -0.394 | P121A | Pro | 121 | Ala | -0.038 | -0.132 |
| L105P | Leu | 105 | Pro | -0.482 | -0.390 | A67V | Ala | 67 | Val | -0.014 | -0.130 |
| V115G | Val | 115 | Gly | -0.314 | -0.383 | A58T | Ala | 58 | Thr | 0.075 | -0.126 |
| M132K | Met | 132 | Lys | -0.525 | -0.370 | G125R | Gly | 125 | Arg | -0.150 | -0.118 |
| F44L | Phe | 44 | Leu | -0.023 | -0.361 | G154V | Gly | 154 | Val | -0.011 | -0.118 |
| V29A | Val | 29 | Ala | -0.119 | -0.361 | V69M | Val | 69 | Met | 0.063 | -0.117 |
| L136Q | Leu | 136 | Gln | -0.502 | -0.356 | K88E | Lys | 88 | Glu | 0.005 | -0.114 |
| I100N | Ile | 100 | Asn | -0.238 | -0.354 | N133D | Asn | 133 | Asp | 0.052 | -0.111 |
| G66S | Gly | 66 | Ser | -0.296 | -0.353 | P89T | Pro | 89 | Thr | 0.119 | -0.111 |
| E7K | Glu | 7 | Lys | -0.370 | -0.350 | R140Q | Arg | 140 | Gln | -0.021 | -0.110 |
| H94R | His | 94 | Arg | -0.266 | -0.346 | V14L | Val | 14 | Leu | 0.020 | -0.110 |
| H83Q | His | 83 | Gln | -0.102 | -0.346 | D127G | Asp | 127 | Gly | -0.067 | -0.109 |
| D142N | Asp | 142 | Asn | 0.044 | -0.346 | A72T | Ala | 72 | Thr | 0.062 | -0.105 |
| G81R | Gly | 81 | Arg | -0.246 | -0.326 | H49Y | His | 49 | Tyr | -0.006 | -0.105 |
| I100F | Ile | 100 | Phe | -0.140 | -0.325 | L150P | Leu | 150 | Pro | -0.085 | -0.105 |
| P101L | Pro | 101 | Leu | -0.084 | -0.324 | A95S | Ala | 95 | Ser | 0.045 | -0.105 |
| A131T | Ala | 131 | Thr | -0.242 | -0.318 | S145C | Ser | 145 | Cys | -0.043 | -0.097 |
| P38A | Pro | 38 | Ala | -0.138 | -0.316 | D127N | Asp | 127 | Asn | -0.044 | -0.096 |
| E19K | Glu | 19 | Lys | -0.219 | -0.316 | R32S | Arg | 32 | Ser | -0.006 | -0.095 |
| V115I | Val | 115 | Ile | -0.175 | -0.313 | D54Y | Asp | 54 | Tyr | 0.034 | -0.093 |
| I108V | Ile | 108 | Val | 0.027 | -0.312 | V18L | Val | 18 | Leu | 0.017 | -0.092 |
| K80N | Lys | 80 | Asn | 0.022 | -0.310 | M143I | Met | 143 | Ile | 0.007 | -0.089 |
| H83R | His | 83 | Arg | -0.019 | -0.308 | A20T | Ala | 20 | Thr | 0.043 | -0.085 |
| G74D | Gly | 74 | Asp | -0.199 | -0.307 | A20V | Ala | 20 | Val | -0.001 | -0.081 |
| A131V | Ala | 131 | Val | -0.142 | -0.306 | I113V | Ile | 113 | Val | -0.011 | -0.079 |
| W8R | Trp | 8 | Arg | -0.057 | -0.303 | A128V | Ala | 128 | Val | 0.042 | -0.078 |
| D21N | Asp | 21 | Asn | -0.190 | -0.302 | G130W | Gly | 130 | Trp | 0.001 | -0.077 |
| G26E | Gly | 26 | Glu | -0.453 | -0.300 | E86D | Glu | 86 | Asp | 0.035 | -0.077 |
| M143T | Met | 143 | Thr | -0.035 | -0.298 | W15L | Trp | 15 | Leu | -0.358 | -0.076 |
| D5Y | Asp | 5 | Tyr | -0.046 | -0.294 | E60D | Glu | 60 | Asp | 0.047 | -0.073 |
| K134N | Lys | 134 | Asn | -0.095 | -0.281 | E137Q | Glu | 137 | Gln | -0.034 | -0.072 |
| A91V | Ala | 91 | Val | -0.090 | -0.274 | P89S | Pro | 89 | Ser | 0.100 | -0.071 |
| G2R | Gly | 2 | Arg | -0.092 | -0.271 | K148N | Lys | 148 | Asn | 0.055 | -0.070 |
| A91G | Ala | 91 | Gly | -0.018 | -0.268 | G125S | Gly | 125 | Ser | 0.028 | -0.067 |
| A95V | Ala | 95 | Val | 0.002 | -0.265 | A144S | Ala | 144 | Ser | 0.049 | -0.066 |
| F34L | Phe | 34 | Leu | -0.014 | -0.261 | H120Q | His | 120 | Gln | -0.220 | -0.063 |
| Y104H | Tyr | 104 | His | -0.044 | -0.260 | G154D | Gly | 154 | Asp | 0.161 | -0.059 |
| H25Q | His | 25 | Gln | -0.347 | -0.260 | G36V | Gly | 36 | Val | 0.032 | -0.058 |
| V11L | Val | 11 | Leu | -0.114 | -0.259 | D45E | Asp | 45 | Glu | 0.047 | -0.056 |
| H120N | His | 120 | Asn | 0.001 | -0.258 | E84G | Glu | 84 | Gly | 0.030 | -0.043 |
| K99T | Lys | 99 | Thr | -0.027 | -0.258 | K51T | Lys | 51 | Thr | 0.061 | -0.043 |
| S4R | Ser | 4 | Arg | -0.130 | -0.257 | R32K | Arg | 32 | Lys | -0.032 | -0.036 |
| S93L | Ser | 93 | Leu | 0.033 | -0.250 | A128T | Ala | 128 | Thr | 0.043 | -0.031 |
| S118G | Ser | 118 | Gly | -0.168 | -0.249 | E106D | Glu | 106 | Asp | 0.019 | -0.028 |
| I76S | Ile | 76 | Ser | -0.328 | -0.244 | D45N | Asp | 45 | Asn | -0.030 | -0.027 |
| K46T | Lys | 46 | Thr | 0.040 | -0.244 | K99N | Lys | 99 | Asn | -0.017 | -0.025 |
| I108F | Ile | 108 | Phe | 0.054 | -0.243 | G125D | Gly | 125 | Asp | -0.006 | -0.025 |
| V115A | Val | 115 | Ala | -0.106 | -0.242 | N13K | Asn | 13 | Lys | 0.013 | -0.021 |
| K17T | Lys | 17 | Thr | -0.073 | -0.242 | G122R | Gly | 122 | Arg | 0.044 | -0.020 |
| V14F | Val | 14 | Phe | -0.019 | -0.230 | N146S | Asn | 146 | Ser | 0.016 | -0.018 |
| D123G | Asp | 123 | Gly | -0.070 | -0.230 | C111G | Cys | 111 | Gly | -0.071 | -0.013 |
| D61N | Asp | 61 | Asn | -0.010 | -0.229 | E137K | Glu | 137 | Lys | -0.124 | -0.006 |
| K79N | Lys | 79 | Asn | -0.088 | -0.224 | H49N | His | 49 | Asn | 0.027 | -0.003 |
| A58V | Ala | 58 | Val | -0.009 | -0.205 | G154S | Gly | 154 | Ser | -0.165 | -0.001 |
| G6R | Gly | 6 | Arg | 0.043 | -0.203 | K148R | Lys | 148 | Arg | 0.049 | 0.000 |
| K97E | Lys | 97 | Glu | 0.015 | -0.202 | E137G | Glu | 137 | Gly | 0.011 | 0.004 |
| E149D | Glu | 149 | Asp | 0.002 | -0.197 | K63R | Lys | 63 | Arg | 0.014 | 0.010 |
| F139L | Phe | 139 | Leu | -0.018 | -0.196 | V14I | Val | 14 | Ile | -0.028 | 0.031 |
| R140W | Arg | 140 | Trp | -0.010 | -0.195 | P23L | Pro | 23 | Leu | 0.080 | 0.033 |
| D5G | Asp | 5 | Gly | -0.051 | -0.194 | G130A | Gly | 130 | Ala | 0.042 | 0.034 |
| G24V | Gly | 24 | Val | -0.027 | -0.193 | Q153H | Gln | 153 | His | -0.019 | 0.035 |
| G2W | Gly | 2 | Trp | -0.169 | -0.184 | V102M | Val | 102 | Met | 0.019 | 0.038 |
| P121L | Pro | 121 | Leu | -0.085 | -0.184 | D45H | Asp | 45 | His | 0.042 | 0.047 |
| K17N | Lys | 17 | Asn | -0.086 | -0.181 | Q153R | Gln | 153 | Arg | -0.058 | 0.058 |
| K57Q | Lys | 57 | Gln | 0.022 | -0.177 | T71N | Thr | 71 | Asn | -0.023 | 0.059 |
| P121S | Pro | 121 | Ser | 0.010 | -0.161 | G130V | Gly | 130 | Val | 0.051 | 0.075 |
| V29I | Val | 29 | Ile | 0.080 | -0.154 | I100T | Ile | 100 | Thr | 0.034 | 0.105 |
| K141E | Lys | 141 | Glu | 0.028 | -0.153 | G6E | Gly | 6 | Glu | 0.108 | 0.111 |
| K119N | Lys | 119 | Asn | -0.576 | -0.150 | T96I | Thr | 96 | Ile | 0.046 | 0.139 |
| H49L | His | 49 | Leu | -0.002 | -0.149 | G75R | Gly | 75 | Arg | 0.004 | 0.164 |
| I31V | Ile | 31 | Val | 0.003 | -0.142 | C111S | Cys | 111 | Ser | 0.132 | 0.250 |
| E55K | Glu | 55 | Lys | -0.125 | -0.140 | | | | | | |

**Table S9)** Hyperparameters and Model Configuration.

| Category | Parameter | Value/Setting |
|---|---|---|
| Pre-trained Models | ESM3 Model Identifier | ESM3_OPEN_SMALL (esm3-open-2024-03) |
| | ProtTrans Model Identifier | Rostlab/prot_t5_xl_half_uniref50-enc |
| MLP Architecture | Input Embedding Dimension | 1536 (ESM3) or 1024 (ProtTrans) |
| | Hidden Layer Sizes | [512, 256, 128] |
| | Activation Function | ReLU |
| | Normalization | Batch Normalization (after each linear layer) |
| | Dropout Probability | 0.6 |
| | Output Dimension | 1 (Regression) |
| Training Hyperparameters | Optimizer | AdamW |
| | Learning Rate | 0.0001 (1e-4) |
| | L2 Weight Decay | 1e-5 |
| | Batch Size | 32 |
| | Loss Function | Mean Squared Error (MSE) |
| | Gradient Clipping (Max Norm) | 1.0 |
| | Training Epochs | 500 |
| | Validation Metric | Coefficient of Determination ($R^2$) |

#### Supplementary Figures

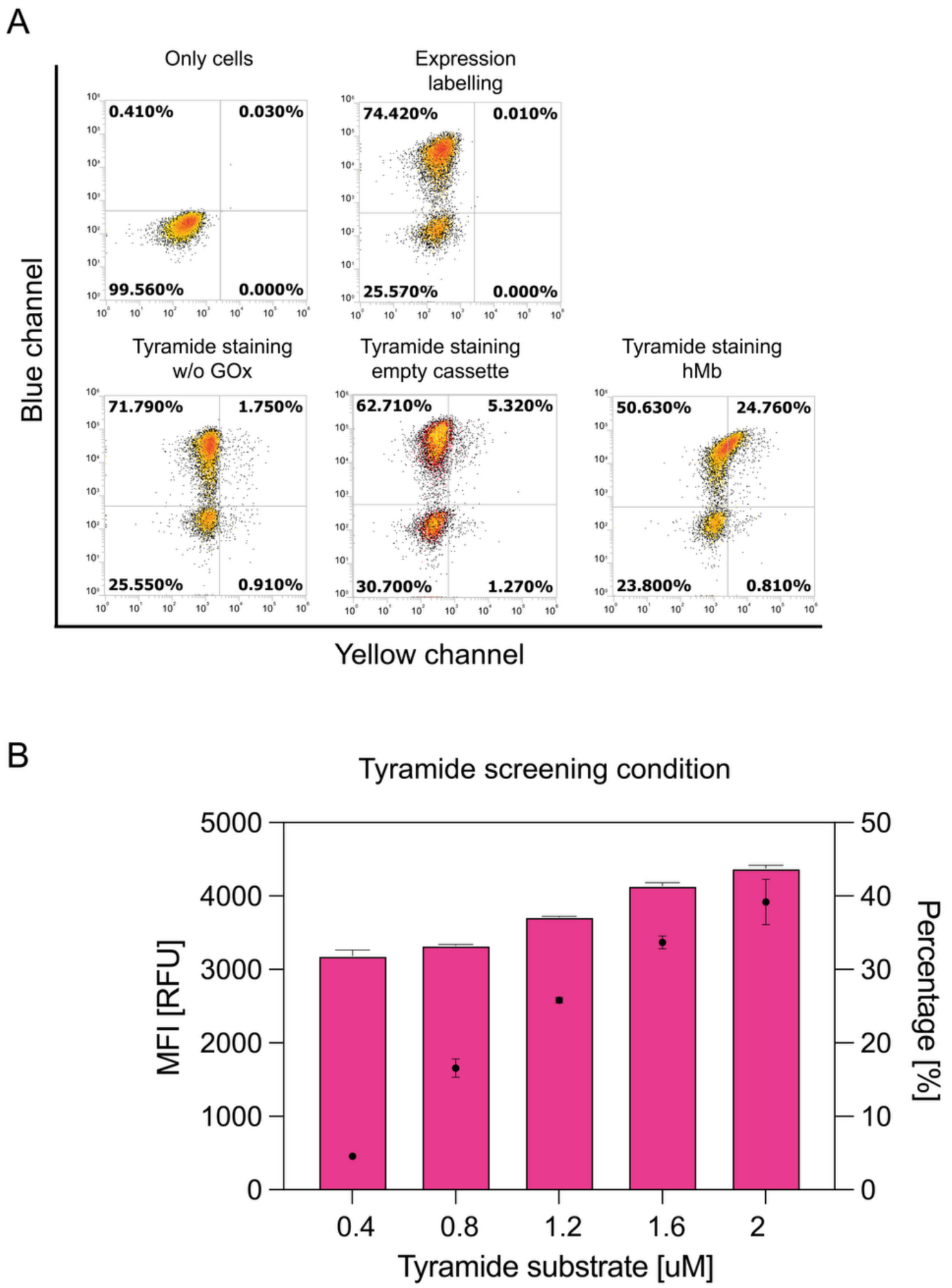


**Figure S1. A)** Yeast tyramide labelling proof of concept, including negative control without glucose oxidase. **B)** Estimation of tyramide AF594 screening compound concentration. Results show mean of median fluorescence intensity (bars) as well as percentage falling in Q1 quadrant for triplicates (scatter). Error bars are standard deviations of triplicates.

A

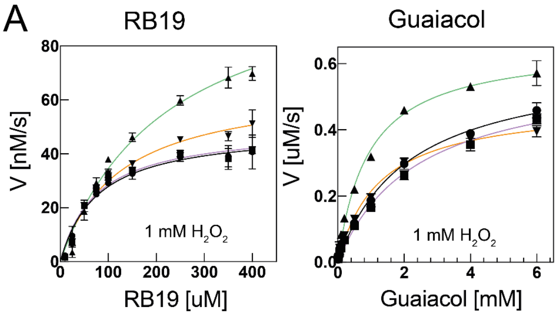


B

WT — Var 4 — Var 9 — Var 14

| | WT | Var4 | Var9 | Var14 |
|---|---|---|---|---|
| Vmax Rb19 [nM/s] | 49.93 ± 3.38 | 50.80 ± 2.90 | 115.63 ± 9.98 | 67.47 ± 5.26 |
| Km Rb19 [uM] | 82.02 ± 16.22 | 83.71 ± 11.15 | 244.36 ± 34.18 | 130.61± 15.31 |
| Vmax Guaiacol [uM/s] | 0.56 ± 0.04 | 0.42 ± 0.05 | 0.64 ± 0.01 | 0.46 ± 0.01 |
| Km Guaiacol [mM] | 9.23± 1.86 | 6.84 ± 1.03 | 4.39 ± 0.19 | 5.85 ± 0.17 |

**Figure S2) A)** Michaelis Menten analysis of machine learning predicted variants along with wild type for guaiacol and reactive blue 19 substrates. **B)** Kinetic constants extracted from figure **A)** Michaelis-Menten fits.

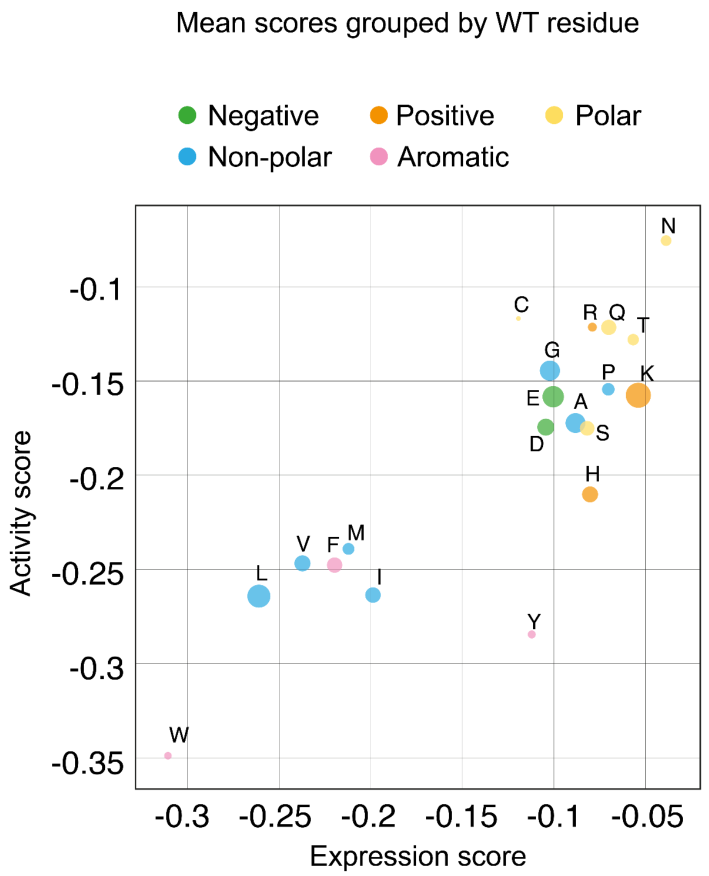


**Figure S3)** Mean expression and activity fitness scores of all single mutations, grouped by WT identity. Sizes of data points correspond to the number of mutations that were averaged. Sample sizes vary across residue types due to uneven representation in the sequence.

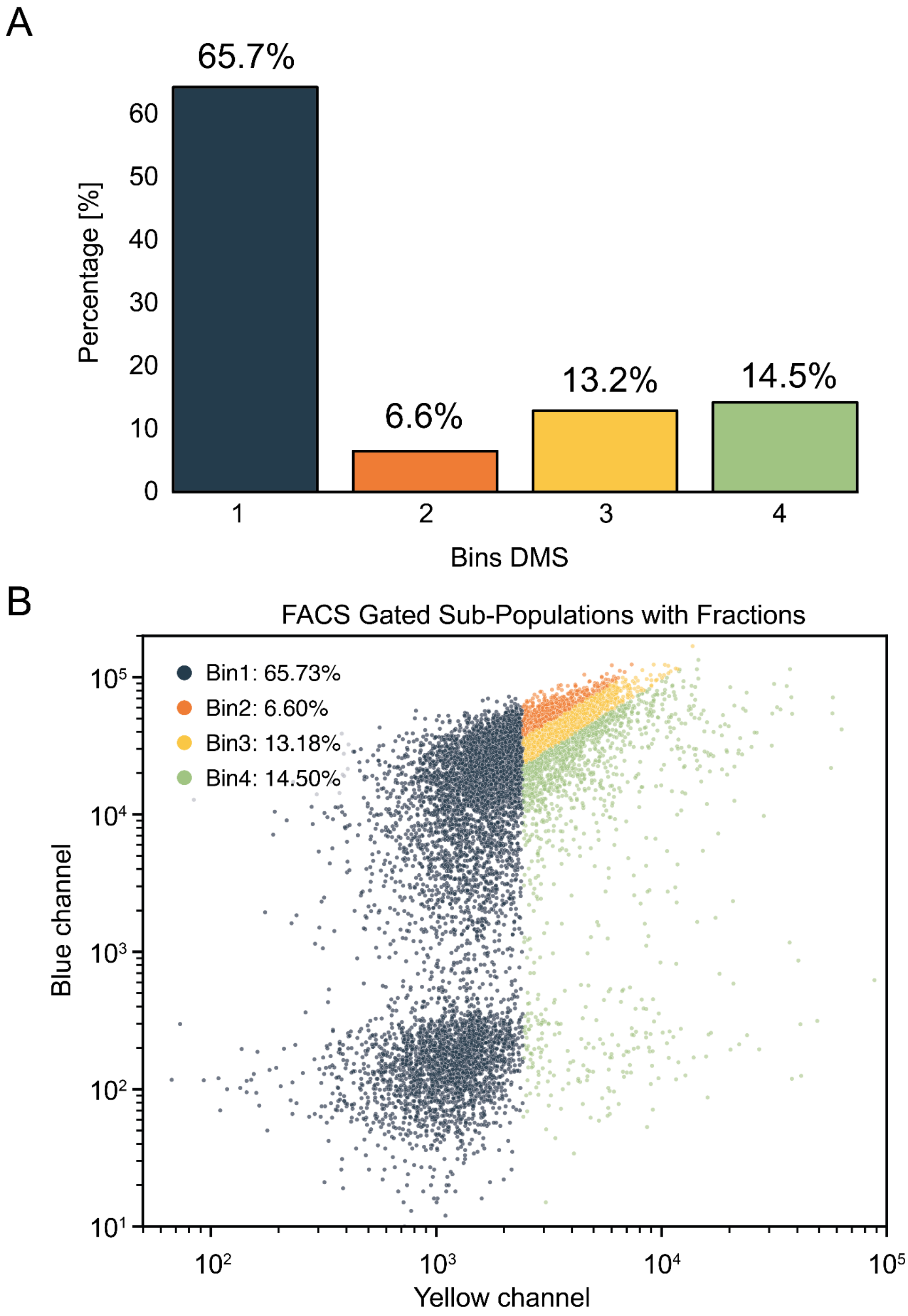


**Figure S4. A)** WT distribution in bins in deep mutational scanning experiment. **B)** Gate settings for monogenic score generation. Gates shown were set to match the WT scores in DMS.

A

| Variant | Pathway | eMap score | EHPath [s] |
|---|---|---|---|
| WT | - | - | - |
| A72W | W72 | 8.64 | $1.39034*10^{-7}$ |
| Q92W | (Y147)- W92 | 13.13 | 0.00583 |
| K97W | W97 | 11.57 | $4.67252*10^{-6}$ |
| F107W | Y104-W107 | 17.00 | $8.52911*10^{-6}$ |

B

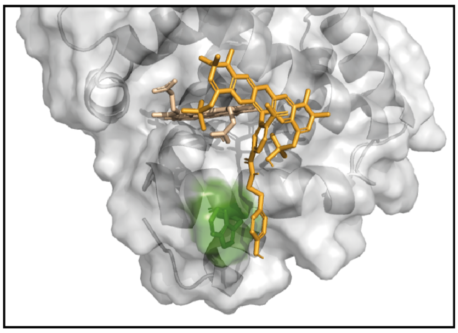

**Figure S5) A)** Table of selected variants and their respective hole hopping paths as suggested by eMap tool as well as the approximate mean residence time as predicted by EHPath.py script[43,44]. The parenthesis for mutant Q92W indicates that while eMap platform suggests direct transfer to W92, EHPaths ranked the transfer through Y147 as being faster - The score and residence time are given for the fastest pathways, respectively. **B)** Molecular docking using Swissdock pose of alphafold generated myoglobin Q92W variant and tyramide. Shown is the 5th best scoring pose, indicating potential pi stacking with the substrate, facilitating electron hopping.

A

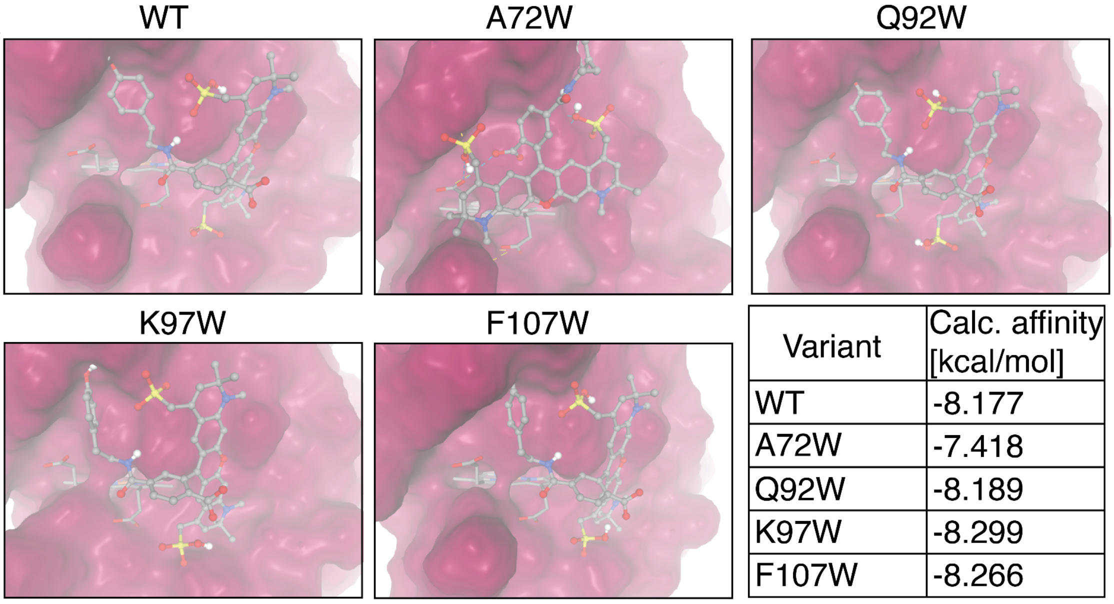


| Variant | Calc. affinity [kcal/mol] |
|---|---|
| WT | -8.177 |
| A72W | -7.418 |
| Q92W | -8.189 |
| K97W | -8.299 |
| F107W | -8.266 |

B

Tyramide AF 594

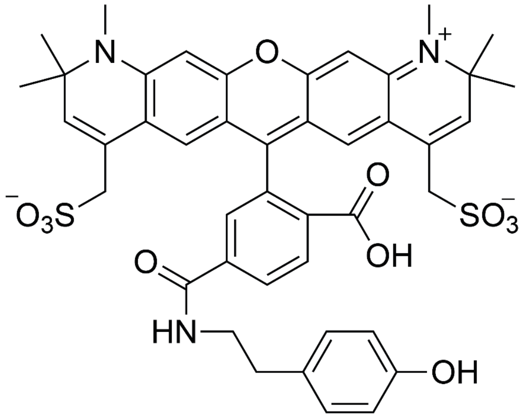


**Figure S6) A)** Docking of tyramide AF594 substrate to mutant variants using Swissdock VINA. Shown is the best ranking pose for all variants. **B)** Chemical structure of Tyramide AF 594.

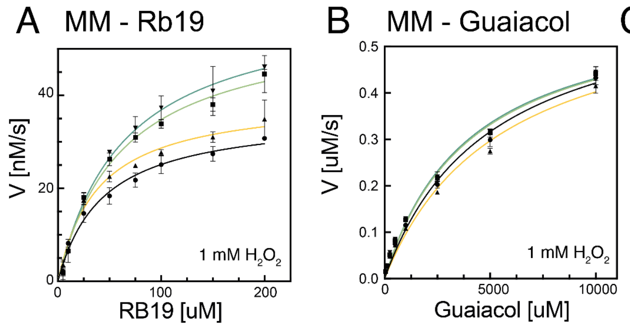


C Kinetic constants

| | WT | Q92W | F107W | Double |
|---|---|---|---|---|
| Vmax Rb19 [nM/s] | 35.34 ± 1.23 | 55.88 ± 4.76 | 39.87 ± 4.79 | 61.93 ± 2.21 |
| Km Rb19 [uM] | 39.75 ± 2.68 | 60.21 ± 6.94 | 38.34 ± 11.62 | 64.36 ± 2.58 |
| Vmax Guaiacol [uM/s] | 0.68 ± 0.05 | 0.64 ± 0.02 | 0.66 ± 0.05 | 0.65 ± 0.02 |
| Km Guaiacol [mM] | 5.00± 0.83 | 3.81 ± 0.76 | 5.31 ± 0.76 | 3.81 ± 0.15 |

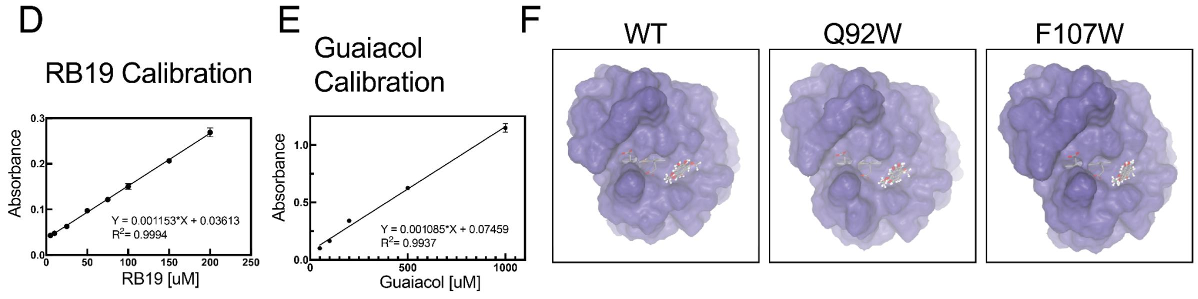


**Figure S7) A) & B)** Michaelis Menten analysis of WT, mutants Q92W, F107W and Q92W/F107W hMb with reactive blue 19 and guaiacol as substrates. Absorbance readout was transformed into molar quantities by calibration curves shown in **D) & E).** Kinetic constants are presented in table in **C)**.
In **F)** we show results of three best docking poses of guaiacol for WT and variants Q92W and F107W. Docking was performed with attracting cavities in Swissdock as described in the method section.

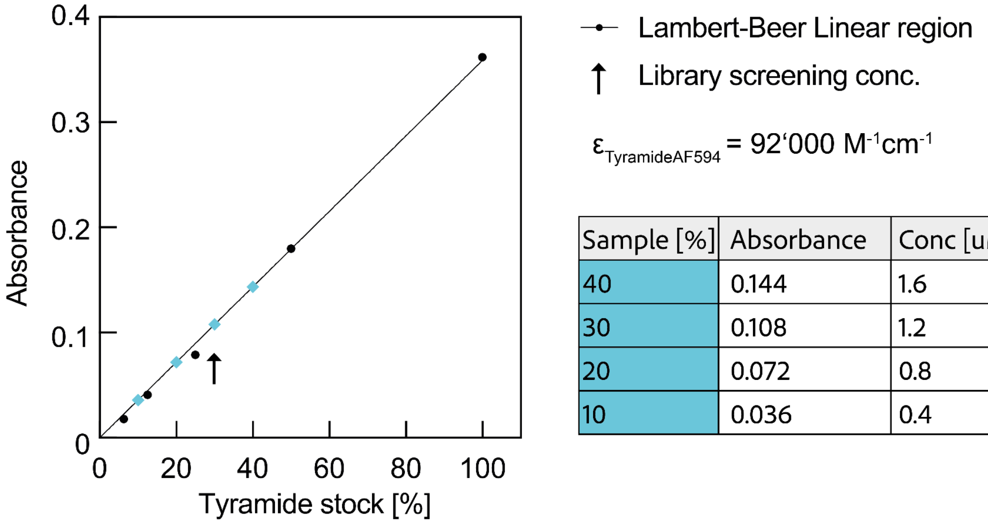


| Sample [%] | Absorbance | Conc [uM] |
|---|---|---|
| 40 | 0.144 | 1.6 |
| 30 | 0.108 | 1.2 |
| 20 | 0.072 | 0.8 |
| 10 | 0.036 | 0.4 |

**Figure S8.** Estimation of concentration of Tyramide AF594. After resuspension in 150 uL DMSO as recommended by the manufacturer, working stocks were produced whereas 100 % corresponds to 1:100 dilution of DMSO stock. Blue data points correspond to concentrations used in Michaelis-Menten experiments given in plots in **Figure 4** and **5**. The black arrow highlights the DMS library screening condition.

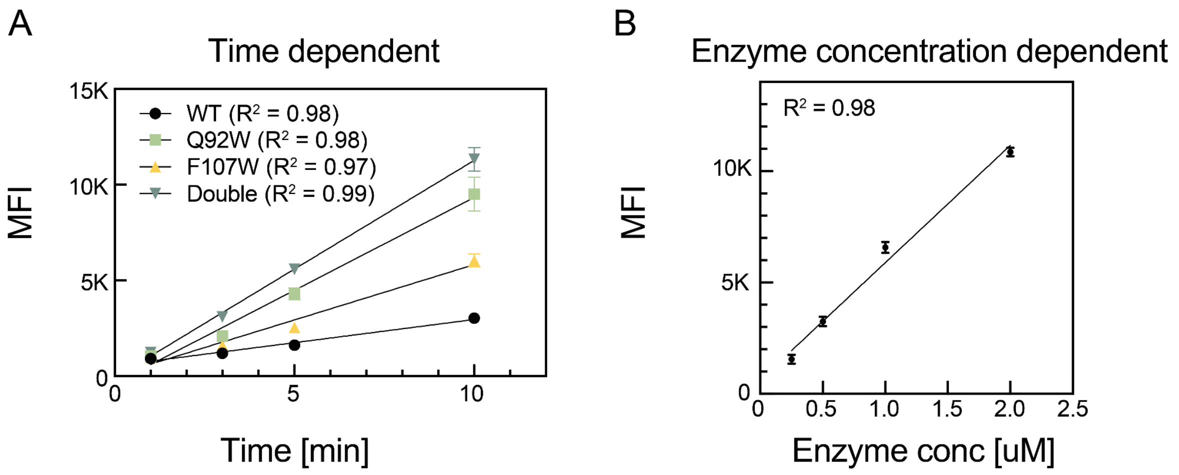


**Figure S9.** Decoupled labelling assay control experiments. Graphs show linearity of MFI of labelled cells over time (**A**) and over enzyme concentration (**B**).
